# Antibiotics select for novel pathways of resistance in biofilms

**DOI:** 10.1101/605212

**Authors:** Eleftheria Trampari, Emma R Holden, Gregory J Wickham, Anuradha Ravi, Filippo Prischi, Leonardo de Oliveira Martins, George M Savva, Vassiliy N. Bavro, Mark A Webber

## Abstract

Most bacteria in nature exist in aggregated communities known as biofilms. Bacteria within biofilms are inherently highly resistant to antibiotics. Current understanding of the evolution and mechanisms of antibiotic resistance is largely derived from work from cells in liquid culture and it is unclear whether biofilms adapt and evolve in response to sub-inhibitory concentrations of drugs. Here we used a biofilm evolution model to show that biofilms of a model food borne pathogen, *Salmonella* Typhimurium rapidly evolve in response to exposure to three clinically important antibiotics. Whilst the model strongly selected for improved biofilm formation in the absence of any drug, once antibiotics were introduced the need to adapt to the drug was more important than the selection for improved biofilm formation. Adaptation to antibiotic stress imposed a marked cost in biofilm formation, particularly evident for populations exposed to cefotaxime and azithromycin. We identified distinct resistance phenotypes in biofilms compared to corresponding planktonic control cultures and characterised new mechanisms of resistance to cefotaxime and azithromycin. Novel substitutions within the multidrug efflux transporter, AcrB were identified and validated as impacting drug export as well as changes in regulators of this efflux system. There were clear fitness costs identified and associated with different evolutionary trajectories. Our results demonstrate that biofilms adapt rapidly to low concentrations of antibiotics and the mechanisms of adaptation are novel. This work will be a starting point for studies to further examine biofilm specific pathways of adaptation which inform future antibiotic use.

## Main

Antimicrobial resistance (AMR) is a complex problem and is a major threat to human and animal health (1). Understanding how bacteria develop resistance to antibiotics is important to inform how they should be used to minimise selection of AMR. There are many genetic mechanisms of antibiotic resistance and the selection of resistant mutants is a classic example of natural selection (2). Initial studies of mechanisms of resistance tended to expose populations to very high concentrations of antibiotics and select for survivors. This identified ‘high-impact’ mutations can confer a large phenotypic benefit and proved very useful for characterising cellular targets and primary resistance mechanisms. However, more recent work has found that repeated exposure to sub-inhibitory concentrations of antimicrobials can have profound impacts on bacterial populations including selection for high level resistance (3,4). This better reflects real world situations where low levels of antimicrobials are commonly present. Importantly, this allows epistatic interactions between multiple genes to be selected and for fitness costs arising from resistance mutations to be ameliorated by additional, compensatory mutations (5).

Much of our understanding of the mechanisms of antibiotic action and resistance comes from laboratory experiments in which bacteria are routinely grown in liquid culture before being exposed to antibiotics. Yet most bacteria in nature exist in biofilms; aggregates of cells often attached to a surface (6). Biofilms represent a fundamentally different mode of life to planktonic cultures and multiple studies have demonstrated extreme changes in gene and protein expression profiles from the same strains when grown in liquid or as a biofilm (7). Many infections include a biofilm component which makes the infection difficult to treat; common examples include infections on prosthetic or indwelling devices. One of the hallmarks of biofilms is inherit resistance to antibiotics, when compared to the corresponding strain grown in liquid culture. One theory explaining the high degree of resistance to antibiotics in biofilms is that cells within a biofilm are metabolically inactive, or even ‘persisters’. In these dormant subpopulations, characterised by arrested macromolecular syntheses, the cellular targets which the antibiotics poison are not essential, thus impeding the bactericidal activity of the antibiotic (8). However, recent studies have demonstrated a strong evolutionary pressure for strains to evolve improved biofilms. In particular, rapid selection of mutants and combinations of mutants with improved biofilm fitness is observed when bacteria are introduced to a new niche (9–11).

Given this evidence of adaptation within biofilms we were interested whether exposure to low concentrations of antibiotics would exert a selective effect on cells within a biofilm. We hypothesised that exposure of biofilms to sub-lethal concentrations of antibiotics would impart a selective pressure and the outcomes of this stress would be distinct to those seen in planktonic cells. To test this hypothesis, we adapted an experimental biofilm evolution model and used *Salmonella* Typhimurium as a model biofilm-forming pathogen which we exposed to three clinically relevant antibiotics. We compared drug exposed biofilm lineages to unexposed biofilm controls and exposed planktonic lineages. We measured the emergence of antibiotic resistance, biofilm capacity, and pathogenicity, and subsequently investigated condition-specific mechanisms of resistance using genome sequencing. We observed rapid adaptation to antibiotic pressure which often carried a cost for biofilm formation, identified novel mechanisms of resistance against cefotaxime and azithromycin and detected biofilm-specific phenotypes showing that studying how biofilms adapt to stress is vital to understanding the evolution of AMR.

## Results

### A biofilm evolution model can study responses to antimicrobial stress

To study the evolution and adaptation of *S.* Typhimurium biofilms when exposed to antibiotics, a serial transfer bead-based model system was adapted and optimised (based on previous work by the Cooper group (11)). To establish *Salmonella* biofilms, we grew bacteria on glass beads in broth (see materials and methods). The beads served as a substrate for biofilms to form on and as a biofilm transfer vehicle, used to move mature biofilms to new tubes with fresh media and new sterile beads. After each transfer, bacteria from the biofilm community had to colonise the new beads and establish biofilms on their surface before being transferred again. This system allows repeated longitudinal exposure of biofilms to the stress of interest and captures all the major components of the biofilm lifecycle. After each passage, cells from biofilms were harvested and populations and individual representative strains were stored and phenotypically characterised. Strains that developed resistance or exhibited altered biofilm formation were selected for whole-genome-sequencing to identify the genetic basis of these phenotypes (Figure 1, a-d).

**Figure 1:**
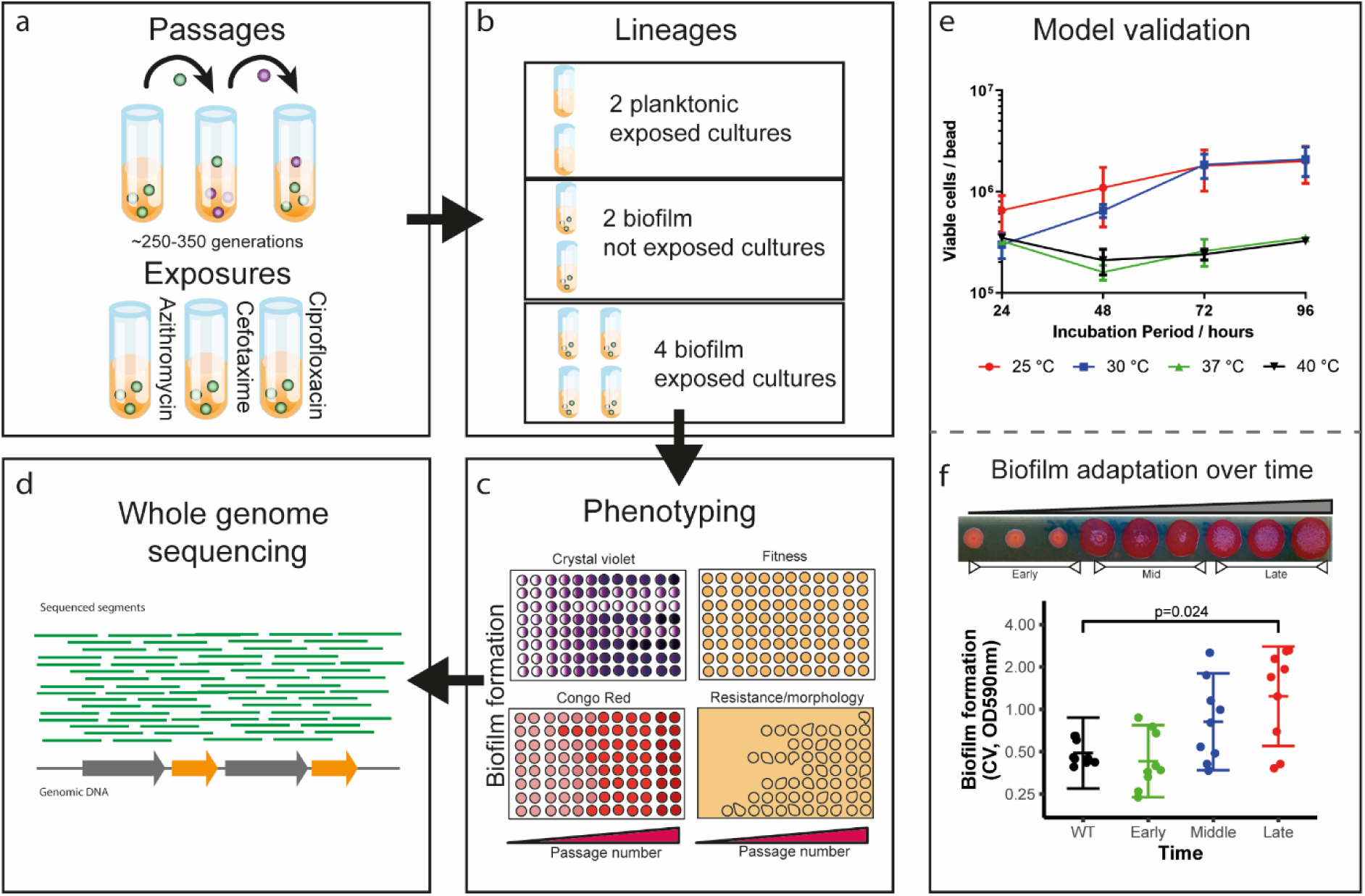
Salmonella biofilm adaptation model. **a,** Sterile glass beads were used for the establishment of Salmonella biofilms. Three antibiotics were selected and used as stressors; azithromycin, cefotaxime and ciprofloxacin. **b,** For each experiment, eight independent lineages were ran in parallel; two planktonic controls, exposed to the drug, with no beads present; two bead controls, not exposed to the drug, and four independent lineages, exposed to the drug on beads. **c,** Passages and sampling were carried out every 72 hours at 30 °C and the cells isolated were stored and phenotyped for biofilm formation, fitness, susceptibility etc. **d,** Over 100 strains (populations and single-cell isolates) were selected and sequenced based on their phenotypic characteristics. **e,** To determine the right experimental conditions for the evolution experiment, cells were recovered and counted from biofilms grown for different periods of time, at different temperatures. This showed that the maximum cell carriage is achieved by 72 h at either 25 °C or 30 °C. Dots indicate average from 4 replicates and error bars show standard error **f.** To determine whether bacteria adapt to the bead model, biofilm formation was monitored over time by visualisation on Congo red-supplemented plates and by the Crystal Violet assay (OD:590nm), using the unexposed bead control lineages. The strains quickly adapted and produced significantly more biomass compared to the WT by the end of the experiment. Each dot represents single cell strains isolated from different timepoints. Error bars reflect estimated +/- one standard error.

To determine the appropriate conditions for the evolution experiments, we measured biofilm formation by *S.* Typhimurium 14028S in lysogeny broth (without salt) after 24, 48, 72 and 96 hours, at 25°C, 30°C, 37°C and 40°C respectively (Figure 1, e). Biofilm formation was determined by measuring cfu per bead (Figure 1, e). Over 72 hours the highest amount of biomass formed (~10^6^ cfu/ bead) was after incubation at 25°C or 30°C; this was stable and consistent. Therefore, we ran the evolution experiments at 30°C with a passage period of 72 hours.

To investigate if and how biofilms would adapt to exposure to sub-inhibitory concentrations of antibiotics, we grew biofilms on beads, in the presence of three clinically-important antibiotics for the treatment of Salmonellosis; azithromycin, cefotaxime and ciprofloxacin. The minimum inhibitory concentrations (MICs) for each antibiotic were determined following the EUCAST microbroth dilution method, adapted to use LB-NaCl and 30 °C to replicate the experimental conditions. Growth kinetics in the presence of all three agents were determined at the same conditions. Based on these results we identified concentrations of each agent (10 μg/mL azithromycin, 0.062 μg/mL cefotaxime and 0.015 μg/mL of ciprofloxacin) that reliably restricted planktonic growth rates to approximately 50% of that of unstressed control cultures which were then used for evolution experiments. This approach had proved tractable in our previous planktonic evolution experiments with biocides (12).

We ran three separate evolution experiments, one with each antibiotic. Within each we included eight lineages; four exposed bead lineages, two exposed planktonic cultures and two drug-free bead-control lineages (Figure 1b). Populations from early, middle and late time points of each experiment were harvested from beads and three single colonies were isolated to allow us to examine phenotypic diversity within the population. The isolates were tested for their biofilm ability, morphology and susceptibility. Biofilm formation was monitored using the Crystal Violet assay (CV) assay and matrix production assessed qualitatively by visualising colonies grown on Congo Red (CR) plates. Susceptibility to antibiotics was measured by determining MICs using the agar dilution method (13).

To confirm the model selects for evolution of increased biofilm formation we phenotyped isolated single cells from drug-free bead-control lineages. We observed an incremental increase in biofilm formation with unexposed isolated colonies forming larger and wrinklier biofilms on CR plates and producing more than three times as much biomass over the course of the experiment (Figure 1, f). This confirmed the model strongly selects for adaptation to produce biofilms with increased biomass over time in the absence of any stressor.

### Biofilms rapidly evolve and adapt in response to sub-inhibitory antibiotic concentrations

To test the phenotypic responses of biofilms repeatedly exposed to non-lethal concentrations of the test antibiotics, we isolated both populations and three individual strains from each lineage at three different timepoints of all experiments; early (first passage), mid (half way point) and late (final passage). We characterised these populations as well as isolated strains for susceptibility to eight clinically-relevant antimicrobials, biofilm formation and colony morphology. We compared results from the biofilm lineages with the corresponding planktonic lineages run at the same time (Figure 2).

**Figure 2:**
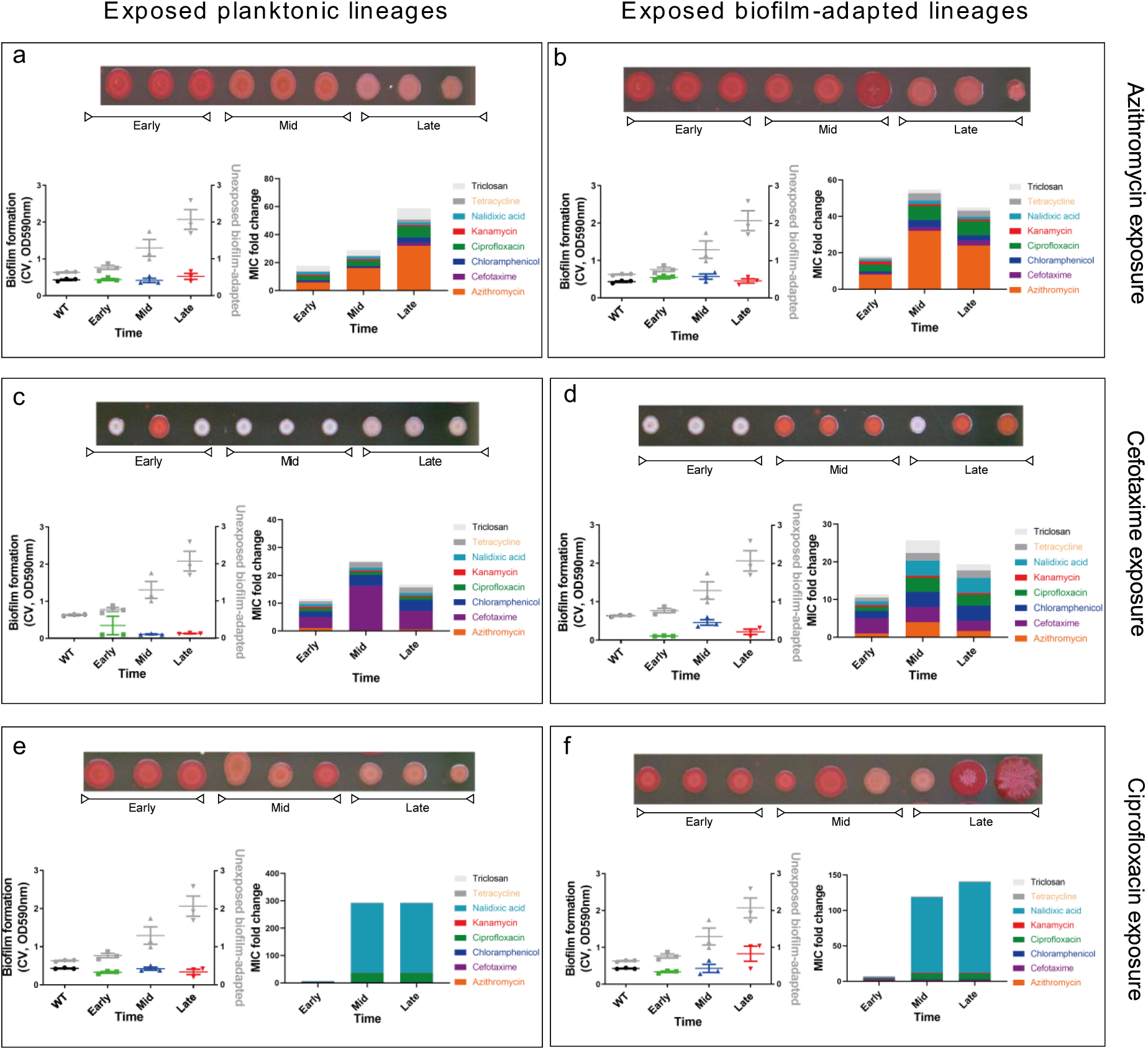
Biofilms adapt to antibiotic stress, with diverse effects on biofilm formation. Planktonic and biofilm populations, exposed to azithromycin (**a-b**), cefotaxime (**c-d**) and ciprofloxacin (**e-f**) were isolated at different timepoints during the evolution experiment (early, mid, late). Panels on the left show data from planktonic lineages, panels on the right from biofilm lineages. Three single isolates per timepoint were tested for their biofilm ability and susceptibility. Biofilm formation was measured by the Crystal Violet assay and on Congo Red plates. Unexposed biofilm lineages were used as a reference to biofilm adaptation (results shown in grey on CV graphs). Antibiotic susceptibility was determined by measuring the MIC values for a panel of different antimicrobials (azithromycin, cefotaxime, chloramphenicol, ciprofloxacin, kanamycin, nalidixic acid, tetracycline and triclosan). Stacked bars were generated by stacking the average MICs for each antibiotic (colour-coded), from three single cell isolates **a-b,** Both planktonic and biofilm isolates, exposed to azithromycin, developed resistance to azithromycin as well as decreased susceptibility to cefotaxime, chloramphenicol, ciprofloxacin, nalidixic acid, tetracycline and triclosan. Biofilm adaptation was inhibited in the biofilm lineages. **c-d,** Planktonic lineages, exposed to cefotaxime, rapidly developed resistance to cefotaxime. Biofilms under the same exposure exhibited an MDR response. Biofilm adaptation of both planktonic and biofilm lineages was completely compromised leading to pale colonies on CR plates. **e-f,** Planktonic and biofilm lineages exposed to ciprofloxacin adapted to the stress by developing resistance to the stressor. Biofilm adaptation was delayed compared to the unexposed control lineages but significantly increased by the end of the experiment.

Biofilms rapidly evolved resistance in response to all three exposures (Figure 2). The time taken to select for emergence of mutants with decreased susceptibility to the antibiotics was similar in both biofilm and planktonic lineages. Azithromycin selected mutants in a stepwise manner with emergence of a population with an 8-fold MIC increase, followed by selection of highly resistant populations with MICs of azithromycin 16 times more than the parent strain. The decreased susceptibility of the azithromycin-exposed lineages became evident at the earliest time point and was fixed by the mid-point of the experiment (Figure 2 a, b). Cefotaxime demonstrated a similar dynamic with both planktonic populations and biofilms exhibiting decreased susceptibility and maintaining this resistance profile until the end of the experiment (Figure 2, c-d). Adaptation to ciprofloxacin resistance was selected by the middle of the experimental period and remained fixed up to the final timepoint in both biofilm and planktonic lineages (Figure 2 e, f).

While the selection dynamics seemed similar between biofilm and planktonic lineages at first glance, analysis of the MICs to all the antibiotics tested, revealed significant differences in the outcomes between biofilm and planktonic conditions. For instance, whilst planktonic populations, exposed to cefotaxime, become mainly resistant to cefotaxime, cefotaxime-exposed biofilms exhibited a multidrug resistance (MDR) phenotype (Figure 2, d). These observations show that whilst the biofilms are able to develop resistance to the selective antibiotics, the mechanisms are likely to be distinct to those seen in planktonic culture.

Whilst it is widely accepted that increased biomass and matrix production improves resilience of biofilms to antimicrobial stress, we observed that biofilm formation itself is heavily influenced by the selective antibiotic. For example, azithromycin prevented the strains from adapting and forming better biofilms whereas unexposed biofilms produced much more biomass over time (Figure 2b). Cefotaxime had a strong negative effect on biofilm formation, with biofilms exposed to cefotaxime actually producing less biomass than the starting wild-type strain and being characterised by pale colony morphology on CR plates (Figure 2d). Ciprofloxacin had less impact on biofilm formation and biofilms exposed to this drug produced increased biomass over time, although to only half the level of the control biofilms. As expected, isolates from planktonic lineages did not form better biofilms over time. On the contrary, cefotaxime exposure again selected for weaker biofilms with a pale colony morphology on CR plates (Figure 2c).

### Individual antibiotics select specific pathways to resistance in biofilms

To investigate correlations between development of resistance to different antibiotics and biofilm formation after each exposure, we compared fold changes in MIC of antibiotics with fold changes in biofilm formation (Figure 3). Each point on the graphs represents a single isolated strain from each evolution experiment (blue: azithromycin exposure, white: cefotaxime exposure, red: cipro exposure, black: drug free exposed controls). All results were compared to averaged data from the parent strain (represented as point ‘0,0’ on the graphs).

**Figure 3:**
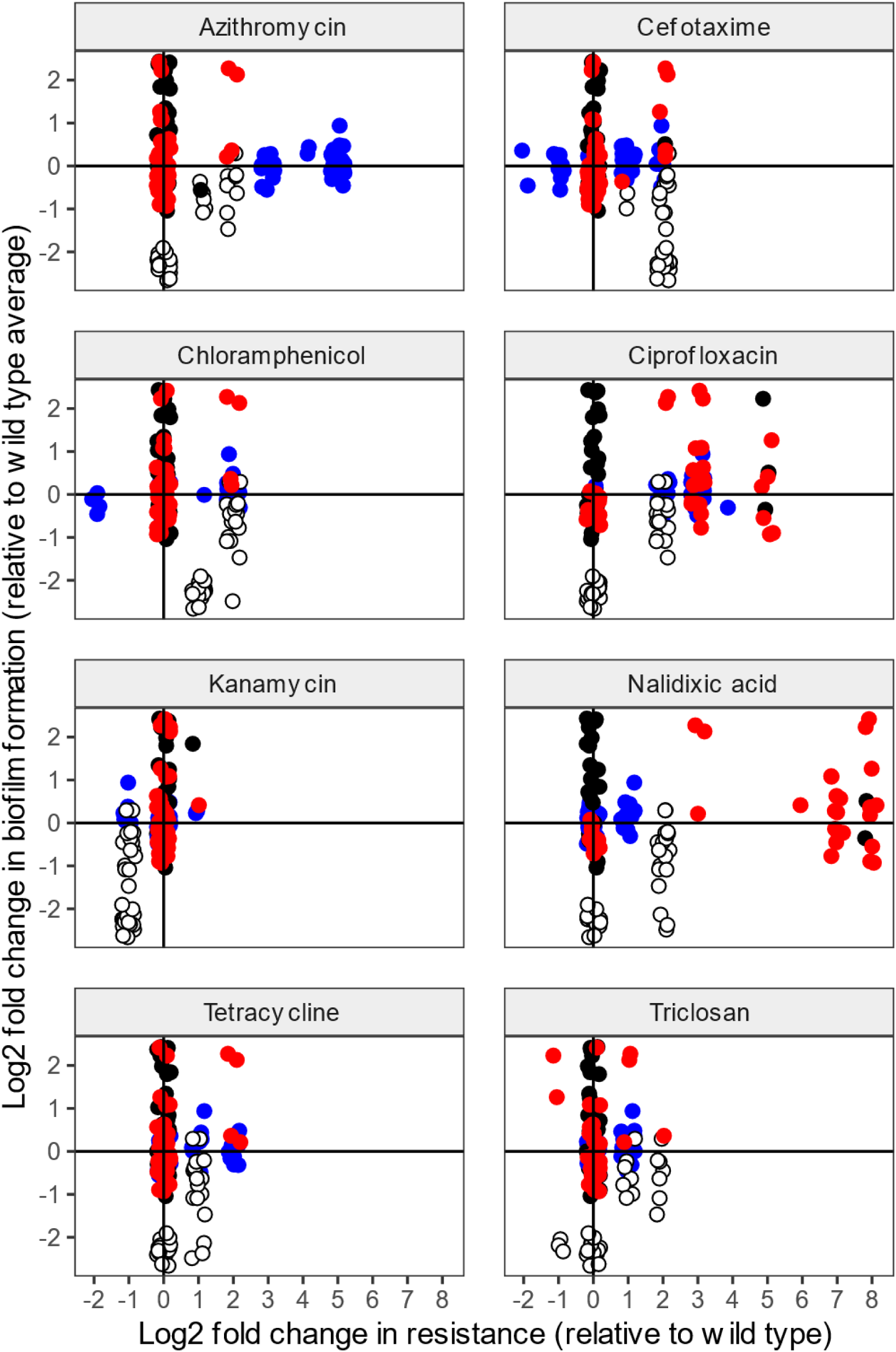
Correlations between resistance and biofilm formation selected by different antibiotics. **a-h,** Fold change in MIC for each antibiotic was compared against the fold change in biofilm formation. Single isolates were characterised; with each dot on the graphs representing an individual isolate from each evolution experiment (blue: azithromycin, white: cefotaxime, red: ciprofloxacin, black: drug-free controls). The parent strain average, used as a reference, is represented as point ‘0,0’ on the graphs. Azithromycin and cefotaxime exposed isolates became less susceptible not only to the stressor but also to other antimicrobials. Ciprofloxacin-exposed isolates only became resistant to fluoroquinolones. Biofilm formation was heavily compromised in cefotaxime-exposed strains. Azithromycin inhibited biofilm formation with some strains exhibiting slightly reduced levels of biofilm formation, whereas ciprofloxacin did not have a significant effect on biofilm adaptation.

As expected, strains became resistant to the antibiotic they were exposed to. There were also examples of selection of cross-resistance to other antibiotics in the biofilms. For instance, azithromycin-exposed isolates exhibited an MDR phenotype and demonstrated decreased susceptibility not only to azithromycin, but also to cefotaxime, chloramphenicol, ciprofloxacin, nalidixic acid, tetracycline and triclosan. Similarly, the cefotaxime-exposed lineages showed increased MICs of chloramphenicol and tetracycline. Most of the antimicrobials tested are known substrates of multidrug efflux pumps (e.g. AcrAB-TolC) except for kanamycin. Strikingly, this was the only antibiotic to which no cross-resistance was observed. In fact, cefotaxime exposed isolates became more susceptible to kanamycin than the parent strains.

Apart from impacts on antibiotic resistance, antibiotic exposure also resulted in major changes in the strains’ ability to form biofilms. Bacteria exposed to cefotaxime exhibited severely compromised biofilm ability, whereas azithromycin and ciprofloxacin led to inhibition or delayed biofilm adaptation respectively. This analysis again indicates that exposure to antibiotics prompts selection for resistance even at sub-inhibitory concentrations, but that this comes at a cost to the ability to form biofilms.

To identify the mechanisms responsible for the phenotypes described above, we whole-genome sequenced over 100 strains (selected to represent major phenotypes of interest and to cover different times in the exposure series for each drug) and identified changes against the parent strain genome. We sequenced a mixture of populations and single cells and used data from single-cell strains to analyse the phylogeny of the mutants (Figure 4a). The results indicated that the different antibiotics selected for mutants which followed distinct paths of adaptation. There was little commonality between the drug exposures showing that there is no universal or generic mechanism of resistance selected for in biofilms. Biofilms exposed to azithromycin and cefotaxime followed a reproducible and distinct evolution pattern (Figure 4b, c), whereas in the ciprofloxacin exposure, bacteria responded to the stress in a number of different ways (Figure 4d). For all drugs, there was separation of biofilm (darker coloured dots) and planktonic lineages (lighter coloured dots) apparent in the phylogeny, again demonstrating different trajectories of selection.

**Figure 4.**
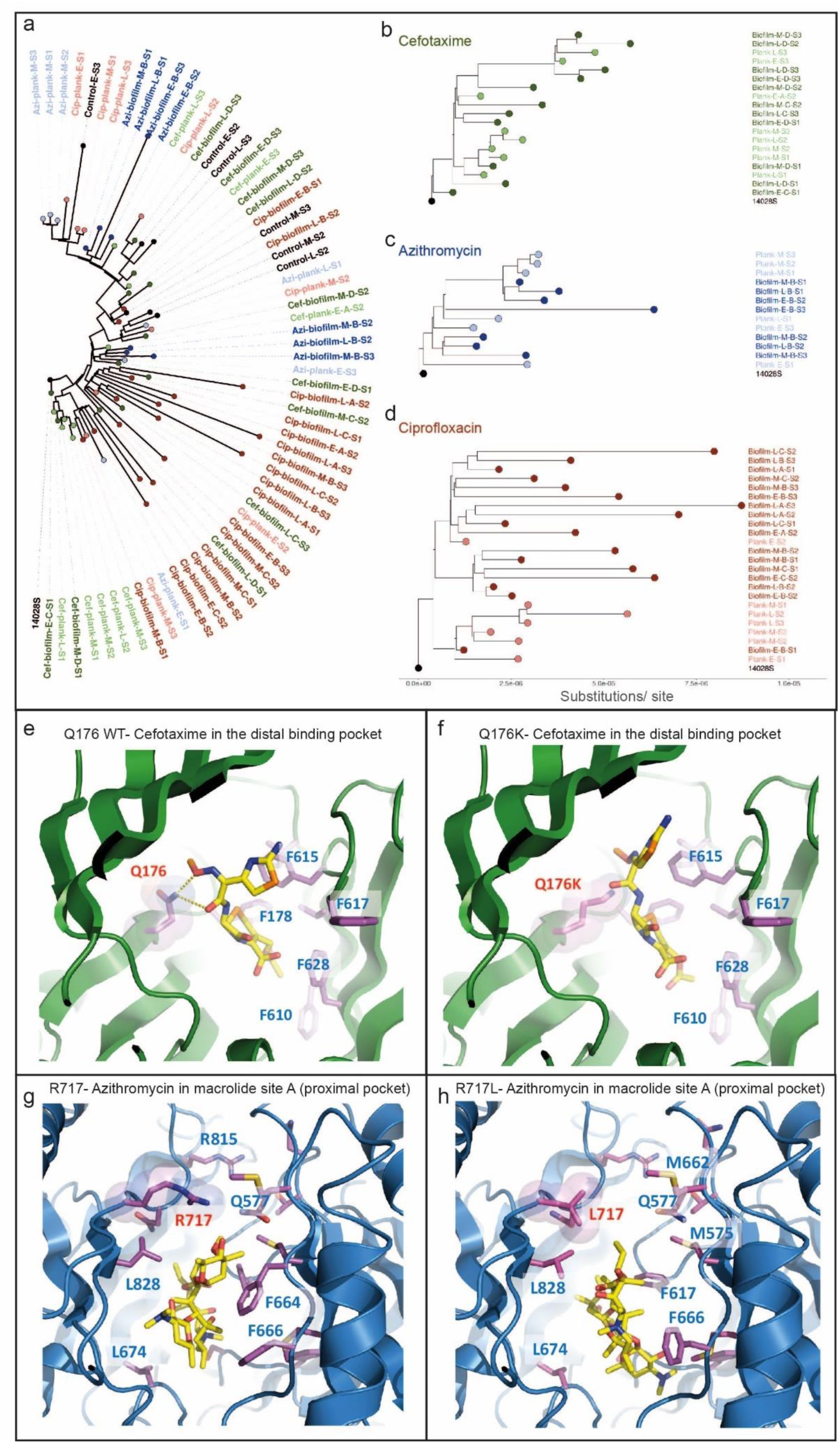
Phylogenetic analysis and AcrB modelling. **a,** Universal phylogenetic tree based on full genome alignment, showing diversity of strains exposed to azithromycin, cefotaxime and ciprofloxacin. Azithromycin and cefotaxime selected for mutants that followed a distinct evolution pattern, whereas ciprofloxacin-exposed strains evolved and responded to the stress in various ways. S. Typhimurium 14028S (CP001363) was used as the reference strain and the tree was arbitrarily rooted at the cultivated parental sequence 14028S. **b-d,** Individual trees were generated for cefotaxime, azithromycin and ciprofloxacin-exposed strains. Dark dots indicate biofilm lineages, light dots planktonic lineages. Phylogenetic variations between biofilms and planktonic cultures were observed, indicating unique mechanisms of resistance between the two states. **e,** Cefotaxime binding to the distal binding pocket of AcrB (4DX5, chain B). Q176 is involved in the coordination of cephalosporin molecules in the binding pocket. **f,** Upon substitution with lysine (Q176K), the free energy of binding changed significantly, potentially resulting in reduced residence time of the drug in the pocket and hence, increased efflux **g,** Azithromycin docking to macrolide site A of the proximal binding pocket of AcrB (3AOB, chain C). R717 exhibits a direct involvement in macrolide coordination in the pocket. **h,** Substitution with leucine (R717L) led to radically altered coordination of azithromycin in the pocket.

### Azithromycin and cefotaxime select novel mechanisms of resistance based upon exclusion of drug from target which in turn hinders biofilm formation

We used the whole-genome sequencing data to identify mutations which may account for potential mechanisms of resistance against the antibiotics of interest. These include some known mechanisms of resistance (e.g. *gyrA* mutation renders bacteria resistant to quinolones (14–16)), as well as Single Nucleotide Polymorphisms (SNPs) that have never been linked to antimicrobial resistance before. The sequence data gave us enough information to build solid hypotheses about the genetic basis for novel mechanisms of resistance against cefotaxime and azithromycin, which we then tested experimentally using a combination of assays (Figure 5).

**Figure 5:**
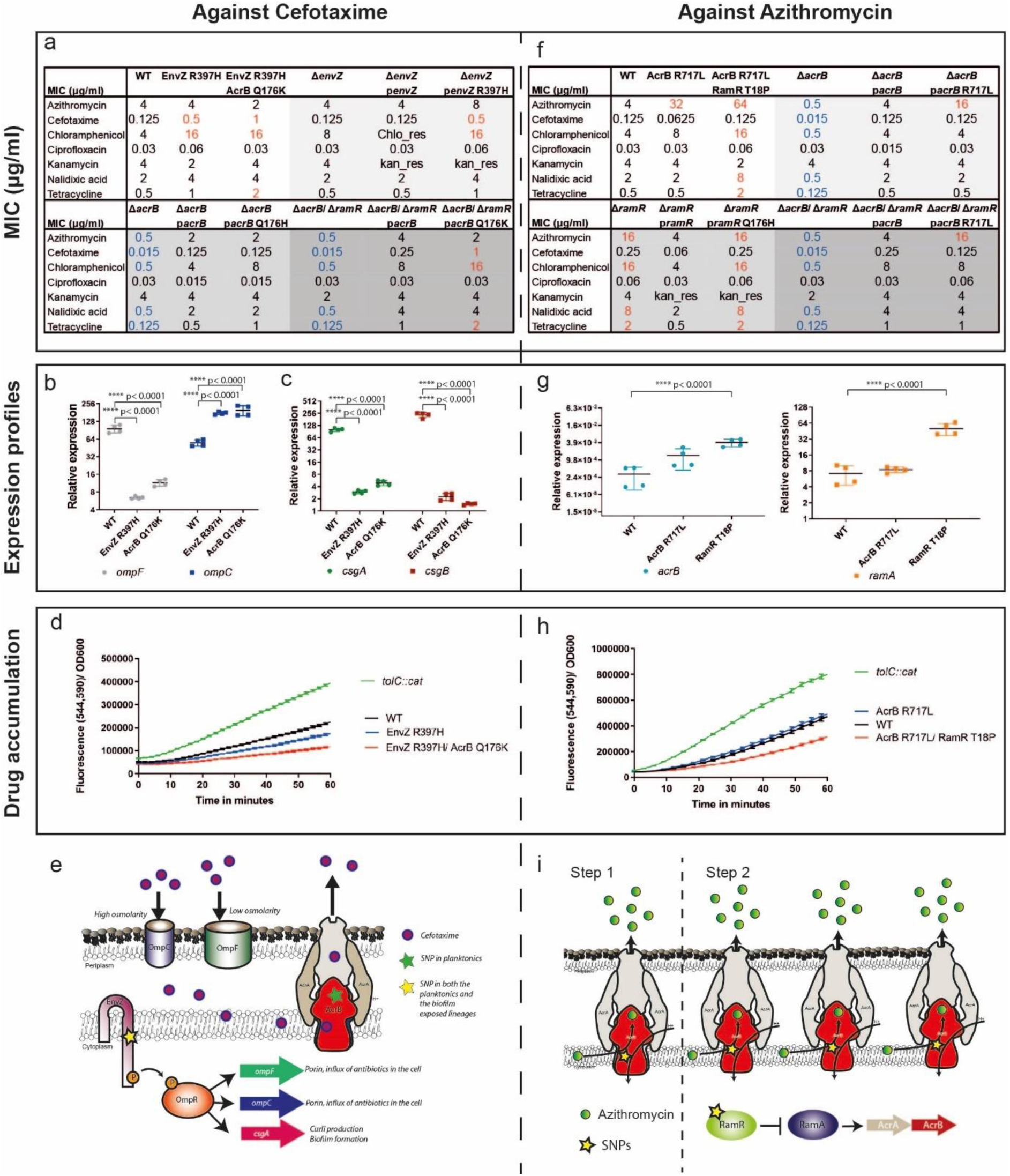
Proposed novel mechanism of resistance to cefotaxime and azithromycin. **a-i,** Cefotaxime and azithromycin exposed isolates exhibiting decreased susceptibility, were whole-genome-sequenced and genetic variations were identified. **a,** Cefotaxime exposed isolates from the middle timepoint carried the R397H substitution in EnvZ leading to reduced susceptibility to cefotaxime and chloramphenicol. Isolates from the late timepoint carried an additional Q176H substitution in AcrB (efflux pump). This resulted in an increase in MIC for both cefotaxime and chloramphenicol as well as tetracycline. Deletion and complementation of WT envZ had no effect on resistance while complementation with the R397H variant, reproduced the resistance results from the isolated strain. Deletion of acrB led to increased susceptibility against all efflux substrate antibiotics. Complementation with AcrB Q176K variant, in the ΔAcrB/ ΔRamR background, resulted in complete recovery of the resistant phenotype. **b,** q RT-PCR from 48-hour biofilms, using gyrB as a reference, resulted in significantly reduced ompF (large porin) expression, whereas ompC (narrow porin) expression significantly increased. **c,** Expression of csgA/B (main curli subunits) was abolished, explaining the pale morphotype cefotaxime-resistant-mutants exhibited. **d,** To test whether the changes on porin composition affect membrane permeability, drug accumulation was measured using the resazurin assay. A tolC::cat, pump-defective mutant, was used as a control. Both resistant mutants exhibited decreased drug accumulation, reflective of the altered membrane composition. **e,** Resistance against cefotaxime is a synergistic result of reduced membrane permeability due to porin alterations and increased efflux through the AcrA/B-TolC pump. **f,** Azithromycin exposed strains, from an early time point, obtained an AcrB R717L substitution which led to an 8-fold increase in azithromycin MIC. At a later stage, an additional substitution in RamR (T18P) emerged. This resulted in an MDR phenotype with increased MICs of azithromycin, chloramphenicol, nalidixic acid and tetracycline. Complementation of AcrB R717L in the ΔAcrB and of the RamR T18P in the ΔRamR background reproduced the resistance profiles of the strains isolated from the evolution experiments, confirming that these substitutions are responsible for the resistant phenotypes observed. **g,** Expression of acrB and ramA were monitored by q RT-PCR in 48-hour biofilms and showed increased expression of both acrB and ramA in the isolate carrying the RamR T18P substitution. **h,** Membrane permeability was monitored by the resazurin assay and reduced accumulation was observed because of the RamR T18P substitution, which potentially leads to overexpression of the RND pump. **i,** Resistance against azithromycin is a result of the modification of the AcrA/B-TolC pump, leading to increased efflux and overexpression of it due to the absence of negative regulation in the RamR T18P substitution strain.

#### Novel mechanisms of cefotaxime resistance

We first observed that isolates that became resistant to cefotaxime had two unique substitutions that they acquired at different timepoints during the evolution experiment. The first substitution was in EnvZ (corresponding to arginine 397 to histidine) and the second was in AcrB (glutamine 176 to lysine).

EnvZ is an osmolarity-sensor protein, attached to the inner membrane of the cell. It functions both as a histidine kinase and a phosphatase, exerting its activity by altering the phosphorylation levels of its cognate transcriptional regulator OmpR. OmpR, amongst other functions, is responsible for differential regulation of the expression of OmpC and OmpF, two principal outer-membrane porins (17,18), as well as curli biosynthesis through the *csgD-csgBAC* operon (19). Mutants carrying the substitution of R397H in EnvZ exhibited a 4-fold increase in the MIC of cefotaxime (Figure 5a) of cefotaxime and produced pale colonies on CR plates. This substitution was present in both planktonic cultures and in biofilms. It is well established that β-lactams cross the outer membrane via porin channels and particularly through OmpF (20–22). It is also well known that OmpR stimulates curli biogenesis by promoting expression of CsgD, which in turn activates the curli operon (23).

AcrB is a key component of the tripartite multidrug efflux pump AcrAB-TolC, and is responsible for substrate recognition and energy transduction (24–26) during the efflux process. Being a member of the *R*esistance *N*odulation cell *D*ivision (RND) family, AcrB shares a common structural organisation with other RND pumps, and couples inward proton transport to antiport (efflux) of a wide range of xenobiotic agents including antibiotics, thus contributing to the emergence of MDR bacteria. The identified Q176K substitution in AcrB was found in strains already containing the EnvZ substitution and led to an additional 4-fold increase in the MIC of cefotaxime (Figure 5a). This substitution was only detected from planktonic cultures and had no additional impact on biofilm formation over that of the EnvZ R397H mutant.

Based on the above, we hypothesised that the ‘first-step’ changes in MIC were due to alterations in expression of *ompC* and *ompF* following mutation of *envZ*. Subsequent substitution in AcrB, was responsible for altered coordination of cefotaxime by AcrB (Q176 is in the distal drug binding pocket of AcrB) and correspondingly lowered residence time of the antibiotic in the pocket, leading to increased efflux of this substrate. Changes in curli expression (due to loss of EnvZ-mediated activation of OmpR), would account for the compromised biofilm formation phenotype we observed for these isolates.

We tested this hypothesis in a number of ways. First, we isolated RNA from 48-hour biofilms, grown in LB agar plates with no salt, formed by strains carrying the identified substitutions (cef-biofilm-M-D-S1 (EnvZ R397H) and cef-plank-L-S2 (EnvZ R397H/ AcrB Q176H)), as well as the WT strain as a control. We performed qRT-PCR to measure *ompC* and *ompF* expression, using *gyrB* expression as our reference (Figure 5b). We observed a significant reduction in *ompF* expression in both the mutants as well as an overexpression of *ompC* indicating that the balance of porin expression had been altered as predicted (Figure 5b). To test whether the decrease in *ompF* led to reduced drug accumulation in the cells, we used a resazurin drug accumulation assay which confirmed that, both mutants show reduced accumulation of drugs inside the cell (Figure 5d).

Additionally, we measured expression of the main curli subunits, *csgA* and *csgB,* and found that expression in the mutants was completely lost (Figure 5c). This explains the pale phenotype on CR and supports our hypothesis that loss of EnvZ function confers protection against cefotaxime, but at a cost to biofilm formation.

We confirmed the specific phenotypic impacts of the two substitutions by creating mutants of the parent strain lacking *acrB* or *envZ* and complementing these with either wild-type or mutant alleles and determining the impacts on phenotypes (Figure 5a). Deletion of *envZ* and complementation with the WT allele did not lead to any MIC changes for any of the antibiotics tested. However, complementation with the mutant pEnvZ R397H allele led to a significant increase in MICs of cefotaxime, chloramphenicol and tetracycline. Similarly, although complementation of the AcrB deletion strain with AcrB or AcrB Q176K did not have an impact on resistance, complementation of AcrB in a ΔAcrB/ ΔRamR background with the Q176K allele led to decreased susceptibility to cefotaxime, chloramphenicol and tetracycline, replicating the phenotype of strains derived from the evolution experiments. These results confirm that the mutations identified are responsible for the resistance phenotypes observed after exposure to cefotaxime.

Finally, we performed *in silico* modelling work to investigate the potential impact of the Q176K substitution on AcrB structure and substrate binding (Figure 4e-f). AutoDock VINA docking simulations were used, allowing for flexible side-chains. Analysis of cefotaxime docking, as well as docking of the control antibiotics, nitrocefin and cephalothin, to both the WT (Q176) and the mutant (Q176K), were based on experimental crystallographic studies (supplementary figure S1, a-b). The analysis focused on the distal binding pocket of chain B of AcrB from *E.coli* (4DX5.pdb) (27) corresponding to the “bound” or “tight” protomer conformation. We found that in both cases, Q176 and the mutant side chain, Q176K, participated in the coordination of cephalosporin molecules. However, the coordination was markedly different, between the WT and the mutant, and resulted in statistically significant reduction in the free energy of binding (ΔG) (Supplementary table S1). This supports the idea that the substitution might cause a reduction in the residence time for drugs in the pocket and as a result, increased efflux and decreased susceptibility.

#### Novel mechanisms of azithromycin resistance

Lineages exposed to azithromycin developed an 8-fold increase in azithromycin MIC (reaching the proposed epidemiological cut-off to define resistance in *Salmonella* (28)) before adapting further to have an MIC 16x higher than the parent (Figure 5f). When we sequenced the resistant strains, we identified two distinct amino acid substitutions. In the first population, there was an arginine 717 substitution to leucine in AcrB and the mutants with highest MICs also had a threonine 18 substitution to proline in RamR.

Interestingly, this substitution in AcrB is distinct from that observed after cefotaxime exposure, and in a different part of the protein. The distal binding pocket is predicted to control access of substrates into the pump, as well as participating in the coordination of high-molecular mass substrates, such as macrolides in the proximal binding pocket (29,30). This indicates that changes in different parts of AcrB confer resistance to different substrates.

RamR is the transcriptional repressor of *ramA*, which is a global transcriptional activator that positively regulates the AcrAB-TolC pump production (31). The inactivation of *ramR* results in over-expression of RamA, consequently increasing pump expression and inducing an MDR phenotype (32). We observed that isolates with this substitution also acquired cross-resistance to cefotaxime, chloramphenicol, nalidixic acid, tetracycline and triclosan, all of which are AcrB substrates.

Our hypothesis was that the R717L substitution in AcrB would result in increased efficiency of efflux of azithromycin and as a result, increased resistance. The additional RamR substitution would then result in over-expression of this mutant pump and explain the further increase in the MIC of azithromycin. To test this hypothesis, we extracted RNA from 48-hour old biofilms and we measured expression of *acrB* and *ramA* by q-RT-PCR, again using *gyrB* expression as our reference (Figure 5g). For both targets there was up-regulation in the mutants compared to the parent strain although expression levels of *acrB* in all cells were low; AcrB is primarily produced in growing cells so low levels of *acrB* mRNA was not unexpected. Intracellular drug accumulation was monitored as described above using the resazurin method (Figure 5h). The R717L mutant alone did not show any changes in accumulation compared to the WT, suggesting the substitution does not impact export of this substrate. This supports the idea that this change improves access for larger substrates to the pump (such as azithromycin, Mr:749), which relies on the proximal drug binding vetting. However, this would not impact a small molecule like resazurin (Mr:229), which has been suggested to directly access the distal binding pocket, by bypassing the proximal pocket of the pump (30,33). Consistent with this idea was the observation of markedly reduced resazurin accumulation in the double mutant, where pump over-expression would be expected to reduce accumulation of a wide range of substrates and confer the typical efflux-based MDR phenotype observed.

To confirm the causative impact of these mutations, we used the same approach as for the cefotaxime mutations where we re-introduced mutant and wild-type alleles of *acrB* and *ramR* into mutants lacking these genes to observe the impact (Figure 5f). AcrB R717L introduction to the *ΔacrB* background led to increased resistance only against azithromycin, complementing perfectly the phenotype of the adapted and evolved strain carrying the AcrB R717L mutation. RamR T18P introduction to the *ΔramR* background led to an additional increase in MICs of azithromycin, chloramphenicol, nalidixic acid and tetracycline, confirming the hypothesis that overexpression of the efflux pump leads to an MDR response. These results confirm that these are the mutations responsible for the resistant phenotypes observed after exposure to azithromycin.

To investigate the impact of the R717L substitution on the structure and function of AcrB, we performed molecular docking of azithromycin and erythromycin (supplementary Figure S2, a-b) in the proximal binding site. The proximal binding pocket is split into two distinct, yet overlapping macrolide binding sites, A and B. Site A is known to be responsible for rifampicin binding, whereas, site B has been associated with erythromycin binding and is located deeper into the proximal pocket (30).

We first docked azithromycin in site B of the proximal binding pocket, producing a model structure, which closely matches the experimental structure of AcrB with erythromycin (3AOC.pbd, chain C, supplementary figure S2, a-b)(30). Our analysis indicated that R717 is located too far away to directly interact with the drug, making it unlikely for the observed coordination in site B to contribute to the observed phenotype. To investigate whether R717 participates in earlier stages of the substrate recognition pathway, we performed docking of azithromycin to site A, located in the front part of the proximal binding pocket (Figure 4g). This showed plausible coordination with direct involvement of R717, and also indicated a clear energetic advantage of the R717 WT over R717L. In fact, R717L led to a radically different coordination of azithromycin (Figure 4h, supplementary table S1), rationalising the observed MDR phenotype.

### Drug resistance in biofilms is stable once selected but costs to virulence and biofilm formation can be ameliorated by stress free passage

We showed that control biofilms that have not been exposed to drugs rapidly adapted to form better biofilms during the experiment (Figure 1f). However, when these strains were tested for drug susceptibility, no changes were observed over time (Figure 6a). There was also no correlation between biofilm ability and antimicrobial susceptibility in these populations which were not exposed to any drugs (Figure 6b). This shows that making more biomass does not alone affect antibiotic susceptibility.

**Figure 6:**
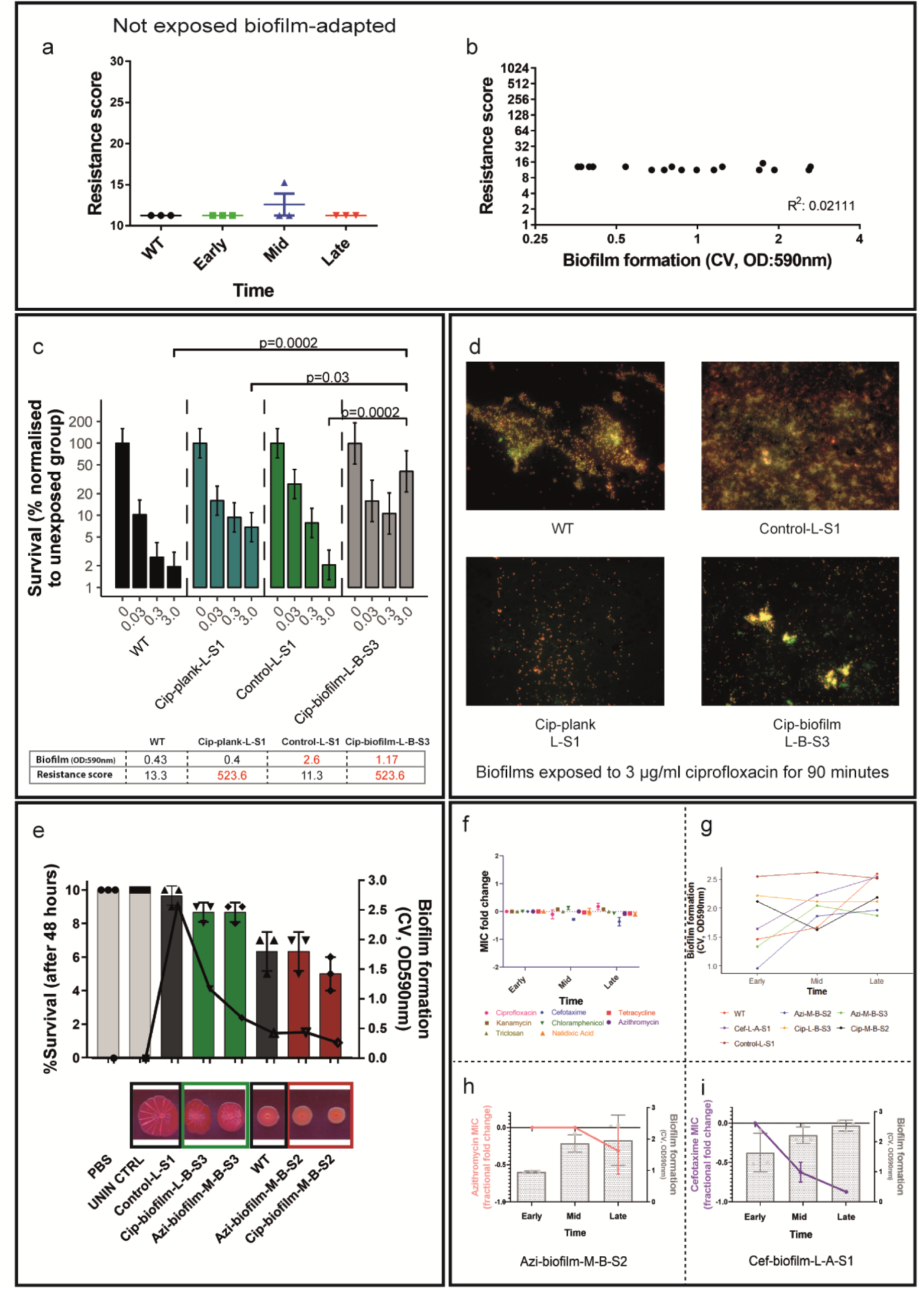
Consequences of resistance. **a,** Unexposed biofilm-adapted lineages were tested for resistance over time with no changes observed in their resistance score (additive value of all MICs determined for a strain). **b,** Although the strains adapted to forming better biofilms over time, their resistance score did not change. **c,** Biofilm viability was tested on 72-hour-biofilms grown on coverslips after treatment with increasing amounts of ciprofloxacin (0, 0.03, 0.3, 3 μg/mL). The strains tested were: cip-plank-L-S1 (resistant but low-biofilm former), control-L-S1 (not resistant but good biofilm former) and cip-biofilm-L-B-S3 (resistant and good biofilm former). Only biofilms produced by cip-biofilm-L-B-S3 were significantly harder to kill with ciprofloxacin. **d,** 72-hour-biofilms grown on coverslips were pre-treated with 3 μg/mL ciprofloxacin for 90 minutes, they were stained with live/dead stain and fluorescence microscopy was performed in a Zeiss Axio Imager M2. Different strains formed biofilms of variable density. An increased number of live cells was only observed in biofilms produced by the cip-biofilm-L-B-S3 strain, with surviving cells forming dense clusters on the coverslip. **e,** Pathogenicity (bars) was tested in the Galleria mellonella infection model. Each point indicates the average number of survivors from independent experiments and the bars show the average of these. The strains tested exhibited resistant phenotypes, with diverse biofilm abilities. WT and the unexposed control-L-S1 strain were used as a reference to the assay. Biofilm formation (lines) was measured by the CV assay and on CR plates. Survival was directly correlated with biofilm formation, with the weak biofilm formers causing more deaths in this model. **f,** Stability of resistance was measured by calculating the average of fold change in MIC per antibiotic, after ten 24-hour passages without any stressor present. For most antibiotics, resistance was stable throughout the accelerated evolution experiment except for cefotaxime. **g,** Biofilm formation increased significantly for the majority of the tested strains, by the end of the experiment. **h,** Individual example of an azithromycin resistant strain, which adapted and formed better biofilm over time without losing resistance to azithromycin. Pink line shows the changes in azithromycin MIC over time (line graph, left axis). Bars (bar chart, right axis) show the average biofilm formation from three replicates. Error bars indicate standard deviation **i,** Individual example of a cefotaxime resistant strain, which adapted to forming better biofilm but lost the resistance to cefotaxime. Purple line shows the changes in cefotaxime MIC over time (line graph, left axis). Bars and error bars are as above.

While we initially investigated whether biofilms have inherently altered susceptibility to antibiotics using conventional susceptibility testing, we also tested whether biofilms were more tolerant of drugs in context. To test whether biofilm-adapted strains have a viability advantage when treated with antibiotics, we grew biofilms on beads for 72 hours at 30°C, and then washed them and exposed them to a range of ciprofloxacin concentrations (Figure 6c). To determine the survival rate, we isolated and counted surviving cells. Survival was normalised against biofilms formed from each strain unexposed to drug. We tested: a planktonic isolate which had been exposed to ciprofloxacin (Cip-plank-L-S1), a control biofilm-adapted isolate not exposed to any drug (biofilm-control-L-S1) and an exposed, biofilm adapted isolate from the ciprofloxacin evolution experiment (Cip-biofilm-L-B-S3). The control biofilms made significantly more biomass than the drug exposed biofilms but demonstrated only a mild increase in survival to the drug treatment compared to the WT strain. Biofilms formed by the drug-exposed planktonic lineage (adapted and highly resistant to ciprofloxacin but with equivalent biofilm formation ability to the parent) had also a mild increase in survival compared to WT biofilms. Interestingly, only bacteria from the biofilm lineage exposed to ciprofloxacin exhibited a significant survival increase even though this strain was no more resistant (by MIC) than the planktonic mutant and made significantly less biofilm than the drug free control. This suggests that neither specific resistance to ciprofloxacin or the ability to make more biomass are enough to improve survival within a biofilm alone. To confirm these results, we grew biofilms on coverslips for 72 hours at 30°C, treated them with 3 μg/mL ciprofloxacin for 90 minutes and carried out live/dead staining to visually characterise survival by fluorescence microscopy (Figure 6d). Our first observation was that, as expected, the different strains formed biofilms of variable density.

Survival was only obvious in the denser biofilm regions, where cells produced more biomass and significant numbers of surviving cells were only observed in the ciprofloxacin-exposed biofilm lineage. Visualisation of this biofilm showed denser clusters of cells than the WT or biofilm-adapted lineage biofilms. Whilst the overall level of biomass was not as high as drug free adapted controls for this strain, the results suggest that a combination of improved biofilm formation and stress adaptation are essential for biofilm viability and survival.

To test whether drug adaptation influences pathogenicity, we selected isolates from different exposures and tested their virulence in the *Galleria mellonella* infection model (Figure 6e, bar chart). Larvae were injected with strains representing different biofilm and resistance phenotypes. Un-injected larvae as well as PBS controls were included as appropriate and the wild type strain (14028S) was used as a reference. In parallel we measured biofilm formation for all cultures used to inoculate larvae using both the CV (Figure 6e, overlaid line graph on the right axis) and the CR assay. We observed that strains with increased biofilm ability were the least infectious, whereas isolates with weaker biofilm phenotypes were most pathogenic. For example, biofilm-control-L-S1, which is a drug free control, biofilm-adapted strain, caused the least deaths with a 95% survival rate. In comparison, Cip-biofilm-M-B-S2, which is a drug-resistant but low-biofilm-forming strain, killed 50% of the larvae. Hence, adaptation to the drug exposure had no obvious advantage for pathogenicity. Our data suggests a strong correlation between pathogenicity and biofilm formation but no association with resistance.

To test the stability of the phenotypes obtained during the course of the evolution experiments, we selected resistant strains to the previous antibiotic exposures with low and high biofilm forming abilities (see materials and methods) and put them through a 24-hour passage, accelerated biofilm evolution experiment (Figure 6, f-i). This was run for 10 days with passages every 24 hours without any antibiotics present, to test whether the resistance and biofilm patterns change over time without any selection. From each population, we isolated single strains and phenotyped them for their biofilm ability and their susceptibility against the same panel of antibiotics we used previously (see materials and methods). We observed that resistance remained stable for most antibiotics (Figure 6f), while the overall ability of the tested strains to form biofilms improved significantly over time (p<0.04, Figure 6g). We also looked individually at the strains with initially low biofilm ability and decreased susceptibility (azi-biofilm-M-B-S2, cef-biofilm-L-A-S1) and we observed that stability of resistance is highly dependent on the nature of the stress. The MIC of azithromycin did not significantly change over time for the azithromycin resistant strain (Figure 6h, orange line), whereas the initially cefotaxime resistant strain exhibited significantly reduced susceptibility by the end of the accelerated experiment (Figure 6i, purple line). In both cases, the biofilm formation of the strains was recovered (Figure 6, h-i).

## Discussion

To develop new antimicrobial strategies, it is important that we understand the genetic mechanisms responsible for bacterial resistance in relevant contexts. Although the evolution of antimicrobial resistance is often studied, previous work has largely focused on planktonic cells and not bacterial biofilms. While it is commonly accepted that biofilms are inherently highly drug resistant, surprisingly little work explores how biofilms evolve in response to antimicrobial stress. One recent report showed that *Acinetobacter* biofilms do adapt to sub-lethal exposure to ciprofloxacin and that mechanisms of resistance were distinct to those seen in planktonic controls (34). Here we address this question using *Salmonella* biofilms as a model system with multiple drugs.

The intrinsic resistance of biofilms to antimicrobials has been attributed to both unique structural characteristics of biofilms but also the wide range of growth states of cells present within biofilms, including dormant or persister cells (35–39). We investigated whether these already resistant communities evolve and adapt further when exposed to sub-inhibitory drug stress. We found that biofilms are highly sensitive to sub-inhibitory exposure to antibiotics, which rapidly selected for changes in the populations. Different stresses selected distinct patterns of adaptation, some of which were unique to the biofilm communities. There was no common response to all three drugs tested, which strongly suggests that there is not a common or generic mechanism for antimicrobial resistance seen in biofilms but instead adaptation highly depends on the nature of the stress.

When cells were exposed to azithromycin or cefotaxime, both planktonic and biofilm populations became resistant not only to the selective drug but also to several different antibiotics, indicating mechanisms conferring MDR. Strikingly, both azithromycin and cefotaxime selected for highly-resistant populations, but also with a marked attenuation of biofilm formation. Lineages exposed to ciprofloxacin were able to improve biofilm formation over time although this was delayed compared to control biofilms. The ciprofloxacin results are consistent with a recent observation from *Acinetobacter* biofilms, where biofilm formation was not compromised after exposure to the drug (34). It is clear from our results that selection of resistance within a biofilm can have a major impact on important phenotypes that impact on bacterial fitness and survival in the real world.

To determine the mechanisms underlying susceptibility, we sequenced more than 100 strains covering the different exposures and time points and demonstrating a range of distinct susceptibility and biofilm phenotypes. Azithromycin exposure selected for the same mutations (changes in AcrB and RamR) in independent lineages, and these emerged in a step-wise manner over the course of the evolution experiment. Similarly, exposure to cefotaxime selected for mutants with altered AcrB and EnvZ, with the same substitution within EnvZ observed in independent lineages. The primary aim of this work was to identify mechanisms rather than the detailed evolutionary dynamics of selection, however the presence of identical substitutions in multiple lineages is strong evidence for the importance of these changes. Ciprofloxacin exposure selected for a wider variety of mutations with much more variation in phenotypes indicating multiple paths of evolution and resistance. The difference between the drugs is likely to reflect the mechanisms of action and resistance; there are multiple known chromosomal mechanisms of ciprofloxacin resistance (including target site changes, porin loss, efflux) whereas high-level resistance to cefotaxime and azithromycin is often a result of acquisition of specific enzymes. In this closed system, acquisition of DNA is not possible, so cells are constrained to mutation of the core genome to generate resistant mutants. This hypothesis was supported by analysis of the mechanisms of resistance.

After analysing the sequencing results, we were able to identify novel mechanisms of resistance against cefotaxime and azithromycin respectively, which we then confirmed experimentally.

We showed that strains exposed to cefotaxime demonstrated a two-step selection of resistance with an initial four-fold rise in cefotaxime MIC, followed by an eight-fold increase. This was linked to an initial R397H substitution in the EnvZ protein followed by an additional Q176K genetic substitution in AcrB. We showed that the EnvZ R397H substitution alters the balance of porin production with down-regulation of the major outer membrane pore, OmpF and up-regulation of the narrow outer membrane pore, OmpC. EnvZ controls activity of OmpR, which is known to control the balance of porin production as well as biofilm formation under low medium osmolarity conditions (17,40). OmpF is well characterised as an entry point for β-lactams due to the molecules’ physiochemical properties, which makes entry possible through this porin which has a larger pore size than OmpC (41,42). We showed that with OmpF expression repressed, the cells are less permeable, leading to reduced levels of the antibiotic in the cell and as a result decreased susceptibility. Whilst the EnvZ substitution was selected in both planktonic and biofilm cultures only planktonic cultures acquired the additional Q176K AcrB substitution, leading to even further reduced drug accumulation and as a result, decreased susceptibility. To date, the exact binding pocket responsible for recognition of cephalosporins, and β-lactams in general, remains debatable with evidence pointing either to the proximal or the distal binding pocket (25,43,44). Here, we showed that Q176K substitution is located in a critically important position within the distal binding pocket of the protein, altering significantly the binding dynamics of the pocket towards cephalosporins. We hypothesise that this may lead to altered residence time in the pocket and result in the observed macroscopic MDR effect. Significantly, our results are consistent with prior data showing that Q176 is involved in substrate transport and coordination of the chromogenic cephalosporin antibiotic nitrocefin in both docking and MD simulation (25,29,45). Intriguingly, the orientation of cefotaxime in the binding pocket was found to differ significantly from that of nitrocefin and cephalothin suggesting multiple modes of β-lactam binding may exist within the distal binding pocket of AcrB (Supplementary Figure S1, a-b, d).

Populations exposed to azithromycin also demonstrated a two-step selection, quickly developing resistance to the drug with an 8-fold followed by a 16-fold increase in azithromycin MIC. These changes were linked to two distinct amino acid substitutions; AcrB R717L and RamR T18P. The AcrB substitution is located in the proximal binding pocket, which is part of the principal drug entry and coordination pathway for high molecular mass drugs and is predicted to impact substrate entry to the pump. A role for R717 in the substrate pathway has previously been proposed (29) and also shown to participate in the coordination of the related antibiotic, rifampicin in at least one experimental structure (30). Docking of azithromycin into the macrolide site A and B of the experimental structures, revealed that the R717 is located far from the canonical site B, which is associated with erythromycin binding. Therefore, this residue is more likely to participate in the earlier stages of the substrate pathway in site A of the proximal pocket. Consistent with this interpretation, we observed that the top docking pose of the azithromycin site A of the proximal pocket brings it in direct contact with the R717 and as a result affected by the R717 substitution. This makes the pump more effective at exporting azithromycin out of the cells and as a result, the population more resistant. The second substitution in RamR resulted in the upregulation of this already “upgraded” AcrAB pump, leading to a 16x rise in azithromycin MIC.

A small number of substitutions within AcrB have been identified and predicted to change affinity for different drugs in the past (46). The substitutions identified in this study have not been previously characterised for their role in resistance, which we did here for the first time, providing strong genetic, phenotypic and structural evidence for their functional impacts.

Both the mechanisms of resistance identified against azithromycin and cefotaxime directly affect both membrane permeability and efflux activity of the cells. The nature of these substitutions leads to cross-resistant phenotypes as accumulation of many drugs is compromised by alterations of general porins and AcrAB. This is of critical importance as exposure to a single drug can select for multi-drug resistant populations with health-threatening implications.

We did however identify clear trade-offs between drug resistance and biofilm formation. Although previous studies have associated exposure to sub-inhibitory concentrations of azithromycin and cefotaxime with inhibition of biofilm formation (47,48), the mechanisms identified in this study have not, to the best of our knowledge, been associated with these antibiotics before. Although we showed that biofilms respond and adapt to antibiotic stresses, we observed that this adaptation is driven by the need to survive exposure to the drug and was not linked to biomass production. Control biofilms passaged without stress made much larger biofilms over time, but these improved biofilm forming lineages did not become more drug resistant. To explore this surprising observation further, we grew biofilms of selected strains representing a variety of biofilm formation and resistance phenotypes and tested their ability to survive exposure to increasing ciprofloxacin concentrations. We observed that only biofilms which had been exposed to ciprofloxacin were significantly harder to kill. This reflects their possession of both a robust community structure and drug-specific resistance mutations that makes them fitter in the specific environment. Neither strains with increased resistance to ciprofloxacin but normal biofilm capacity, nor those with normal drug sensitivity but increased biofilm capacity, demonstrated a significant benefit when treated with ciprofloxacin. Based on these results, we hypothesise that producing more biomass is not necessarily the best solution to survive antibiotic exposure. Highly resistant biofilms may be more likely to result from a combination of both structural and drug specific mechanisms.

Interestingly we did not identify mutations in pathways previously proposed to contribute to persister cell formation, suggesting that these were not important in adaption to the drug exposures in our experimental setup.

Biofilms play a crucial role in chronic infections and our observations suggested an obvious fitness advantage of adapted biofilms over unexposed biofilm populations in terms of drug resistance. To see if this impacts virulence we investigated the pathogenicity of strains with different resistance and biofilm profiles, using the *Galleria mellonella* infection model. We observed that mutations that rendered the bacteria resistant to drugs had no significant impact on pathogenicity. However, the biofilm ability of the strains was negatively correlated with pathogenicity, with strains forming least biofilm being most virulent resulting in the lowest survival rates.

Having characterised a number of biofilm-related resistant phenotypes, we estimated their stability in the absence of drug selective pressure using an accelerated biofilm evolution experiment. Strains that had been exposed to ciprofloxacin and azithromycin maintained their resistance profiles over extended passaging but formed better biofilms. In contrast, cefotaxime exposed populations lost their acquired resistance after a few passages whilst they became better biofilm formers. This indicates that although stability of resistance is highly influenced by the nature of the antimicrobial stress, bacteria can quickly adapt to a more sessile, community-orientated lifestyle in the absence of drug. Analysis of azithromycin-exposed populations which had improved their biofilm ability identified loss-of-function mutations in cyclic di-GMP phosphodiesterase, YjcC. This is unsurprising as cyclic di-GMP is well known for its role in biofilm formation in several organisms including *Salmonella*, which harbours 12 proteins with GGDEF and 14 proteins with EAL domains (49,50).

In conclusion we demonstrate here that biofilms are highly sensitive to stress from low levels of antibiotics, rapidly adapt to drug pressure and that mechanisms of resistance can incur costs to other important phenotypes. Using similar approaches to those outlined here will help understand the impacts of drug exposure on biofilms in many contexts. This can help inform how best to use antimicrobials and predict how biofilms will respond to different stresses.

## Materials and methods

### Biofilm adaptation and evolution model

*Salmonella enterica* serovar Typhimurium 14028S was used as the parent strain to initiate all biofilm experiments in this study. This strain has been used as a model for *S.* Typhimurium biofilm studies by many groups including our own and has a fully closed and annotated reference genome (Accession number: CP001363). To study adaptation and evolution of *Salmonella* biofilms, we adapted a model described by the Cooper group (11). Bacteria were grown on 6 mm soda lime glass beads (Sigma, Z265950-1EA) for 72 hours in Lysogeny Broth (LB) with no salt. They were incubated in glass universal tubes containing 5 mL of the medium in horizontal position, with mild rocking at 40 rpm, at 30 °C. For each passage, the beads were washed in PBS and transferred into fresh media with new sterile beads. The experiment was carried out at the presence of three clinically-important antibiotics; azithromycin, cefotaxime and ciprofloxacin at a final concentration of 10 μg/mL, 0.062 μg/mL and 0.015 μg/mL respectively. Eight independent lineages were included per exposure; four drug-exposed biofilm lineages, two drug-exposed planktonic cultures and two unexposed, bead-only control lineages. In each tube, three initially sterile beads were used, one to be transferred to the next lineage, one to be stored, and one from which cells were recovered for phenotyping. For storage, one bead per passage was frozen in 20 % glycerol. For phenotyping, the cells were isolated from the beads by vortexing in PBS for 30 seconds and then grown overnight in 1 mL of LB broth, before being stored in chronological order in deep-well plates in glycerol. The experiments were completed after 250 generations (17 passages) for the azithromycin and cefotaxime exposure and after 350 generations (24 passages) for the ciprofloxacin exposure. Populations from an early (first passage), middle (half way point lineage) and late (final passage) time point were chosen for study and from each, three single colonies were isolated, sub-cultured and stored in 20% glycerol. These single-cell isolates, and their parent populations were stored in deep-well 96-well plates and used for phenotyping by replicating the bacteria onto appropriate media to test for fitness, biofilm ability, morphology and susceptibility (replication used ‘QRep 96 Pin Replicators’, Molecular devices X5054). Figure 1 shows an overview of the experimental setup and phenotyping procedure.

### Model optimisation

To determine the optimum culture conditions for achieving the greatest cell carriage of *S.* Typhimurium 14028S biofilms on the glass beads, biofilms were grown in 5 mL LB without salt on 6 mm glass beads at four standard microbiological incubation temperatures: 25 °C, 30 °C, 37 °C and 40 °C. The cell counts on beads grown at each temperature was determined every 24 hours for 96 hours. Biofilms were washed in 1 mL PBS and harvested via vortexing for 30 seconds. The harvested cells were serially diluted in a microtiter tray containing 180 µL PBS and 5 µL was spotted onto a square LB agar plate. The number of colony forming units was calculated and the incubation conditions yielding the greatest amount of cells was determined.

### Crystal violet assay (CV)

To measure biofilm formation, selected strains were grown overnight in LB broth and then diluted into 200 μL of LB-NaCl to give an OD of 0.01 in microtiter plates. The plates were incubated at 30 °C for 48 hours, covered in gas-permeable seals before wells were emptied and vigorously rinsed with water before staining. For staining, 200 μL of 0.1% CV was added to each well and incubated for 15 minutes at room temperature. The crystal violet dye was then removed, and the wells were rinsed with water. The dye bound to the cells was then dissolved in 70% ethanol and the absorbance was measured at 590 nm in a plate reader (FLUOStar Omega, BMG Labtech).

### Biofilm morphology

To visually assess biofilms morphology, we replicated isolates stored in 96 deep-well plates on 1% agar LB-NaCl plates, supplemented with 40 μg/mL Congo red (CR) dye. The strains of interest were diluted to a final OD of 0.01 in a microtiter plate and were then printed on the Congo red plates. The plates were incubated for 48 hours at 30 °C before being photographed to capture colony morphology.

### Antimicrobial susceptibility testing

To determine the minimum inhibition concentrations of antimicrobials against strains of interest, we used the broth microdilution method (51) and the agar dilution method (13), following the EUCAST guidelines. In both cases, Mueller-Hinton broth or agar was used. Changes of less than two dilutions were not considered significant.

### Molecular Modelling and Antibiotic Docking

For docking analysis, protein and ligand PDB files were first converted to PDBqt format using Raccoon (52). AutoDock Vina (53) was used for unbiased flexible docking simulations of ligands on protein structures using an optimal box size, estimated according to the radius of giration of the ligands (grid-point spacing of 1 Å), centred on the predicted binding sites (supplementary table S1). The PyMOL Molecular Graphics System, Version 2.0 Schrödinger, LLC. was used to visualize the results. Formore information on model validation see supplementary materials.

### Extraction of DNA

To extract genomic DNA for sequencing, selected strains were grown O/N in a 96-deep-well plate in LB, at a final volume of 1.5 mL. Cells were recovered by centrifugation at 3,500 g and were resuspended in 100 μL of lysis buffer (5 μg/mL lysozyme, 0.1 mg/mL RNAse in Tris-EDTA, pH 8) per well. The resuspended cells were then transferred in a new semi-skirted, low-bind PCR plate, secured with an adhesive seal and incubated at 37 °C, 1600 rpm for 25 minutes. 10 μL of a lysis additive buffer (5% SDS, 1 mg/mL proteinase K, 1 mg/mL RNAse in Tris-EDTA, pH 8) was added in each well and the plate was sealed with PCR strip lids before being incubated at 65 °C, 1600 rpm for 25 minutes. The plate was briefly centrifuged and 100 μL were moved to a new PCR plate. For the DNA isolation, 50 μL of DNA-binding magnetic beads (KAPA Pure beads, Roche diagnostics) were added in each well and were incubated at room temperature for 5 minutes. The plate was then placed on a magnetic base and the supernatant was removed by pipetting. The beads were washed three times with 80% freshly-prepared ethanol. After removing the last wash, the beads were left to dry for 2 minutes before eluting the DNA. For the DNA elution, the plate was removed from the magnetic apparatus and 50 μL of Tris-Cl were added to each well. The beads were pulled using the magnetic apparatus and the isolated DNA was transferred to a new PCR plate. DNA concentration was determined using the Qubit ds DNA HS Assay kit (Q32851) following the manufacturer’s instructions.

### Whole Genome sequencing

Genomic DNA was normalised to 0.5 ng/µL with 10mM Tris-HCl. 0.9 µL of TD Tagment DNA Buffer (Illumina Catalogue No. 15027866) was mixed with 0.09 µL TDE1, Tagment DNA Enzyme (Illumina Catalogue No. 15027865) and 2.01 µL PCR grade water in a master mix and 3ul added to a chilled 96 well plate. 2 µL of normalised DNA (1ng total) was mixed with the 3 µL of the tagmentation mix and heated to 55 ⁰C for 10 minutes in a PCR block. A PCR master mix was made up using 4 ul kapa2G buffer, 0.4 µL dNTP’s, 0.08 µL Polymerase and 4.52 µL PCR grade water, contained in the Kap2G Robust PCR kit (Sigma Catalogue No. KK5005) per sample and 11 µL added to each well need to be used in a 96-well plate. 2 µL of each P7 and P5 of Nextera XT Index Kit v2 index primers (Illumina Catalogue No. FC-131-2001 to 2004) were added to each well. Finally, the 5 µL Tagmentation mix was added and mixed. The PCR was run with 72 ⁰C for 3 minutes, 95 ⁰C for 1 minute, 14 cycles of 95 ⁰C for 10 seconds, 55 ⁰C for 20 seconds and 72 ⁰C for 3 minutes. Following the PCR reaction, the libraries were quantified using the Quant-iT dsDNA Assay Kit, high sensitivity kit (Catalogue No. 10164582) and run on a FLUOstar Optima plate reader. Libraries were pooled following quantification in equal quantities. The final pool was double-spri size selected between 0.5 and 0.7X bead volumes using KAPA Pure Beads (Roche Catalogue No. 07983298001). The final pool was quantified on a Qubit 3.0 instrument and run on a High Sensitivity D1000 ScreenTape (Agilent Catalogue No. 5067-5579) using the Agilent Tapestation 4200 to calculate the final library pool molarity. The pool was run at a final concentration of 1.8 pM on an Illumina Nextseq500 instrument using a Mid Output Flowcell (NSQ® 500 Mid Output KT v2(300 CYS) Illumina Catalogue FC-404-2003) and 15 pM on a Illumina MiSeq instrument. Illumina recommended denaturation and loading recommendations which included a 1% PhiX spike in (PhiX Control v3 Illumina Catalogue FC-110-3001). Whole genome sequencing data has been deposited in the Sequence Read Archive under PRJNA529870.

### Bioinformatics

Sequence reads from the sequencer were uploaded on to virtual machines provided by the MRC CLIMB (Cloud Infrastructure for Microbial Bioinformatics) project using BaseMount (54). Quality filtering of the sequence reads was performed using Trimmomatic (version 3.5) with default parameters (55). Trimmomatic’s Illuminaclip function was used to remove the Illumina adapters. The trimmed reads were then assembled into contigs using SPAdes version 3.11.1 using default parameters (56).

To determine single nucleotide polymorphisms (SNPs) between the de novo assembled Salmonella genomes and the parent genome. Snippy version 3.1 was used using parameters recommended in (https://github.com/tseemann/snippy). The Snippy-core from the Snippy tool box was used to determine the core SNPs. The full genome alignment output by Snippy-core was used in subsequent phylogenetic analyses, after removal of the published reference sequence (accession number CP001363). All 4870267 sites were included in the analysis to avoid ascertainment bias (57). A whole-genome phylogenetic tree was then inferred from the 63 sequences under the model HKY+G implemented in iq-tree (58). For the individual population analyses the control sequences were removed, the trees were estimated with iq-tree, using the HKY+G evolutionary model. All trees were arbitrarily rooted at the cultivated parental sequence 14028S for visualisation purposes, and were plotted with ggtree for R (59). Branch lengths are given in units of substitutions/site.

### Preparation of RNA samples for q-RT PCR

RNA from biofilms was isolated using the SV Total RNA Isolation System kit (Promega). RNA was extracted from the WT (14028S), cef-biofilm-M-D-S1 (EnvZR397H), cef-plank-L-S2 (EnvZR397H/ AcrB Q176K), azi-biofilm-E-B-S2 (AcrB R717L) and azi-biofilm-M-B-S1(AcrB R717L/ RamR T18P) strains. These were grown O/N at 37 °C in LB and then spotted on LB-NaCl agar plates and were grown for 48 hours at 30 °C. Cells from each spot were then resuspended in 100 μL TE containing 50 mg/mL lysozyme and were homogenised by vortexing. 75 μL RNA Lysis Buffer (Promega kit), followed by 350 μL RNA Dilution Buffer (Promega kit) were added to the cell suspensions, which were then mixed by inversion. Samples were incubated at 70 °C for 3 minutes and centrifuged at 13,000 g for 10 minutes. The supernatant was mixed with 200 μL 95 % ethanol and was then loaded on to the spin columns provided by the kit. The columns were washed with 600 μL RNA Wash Solution. DNAse mix was prepared following the Promega kit protocol and 50 μL were directly added on the column membrane. After a 30 minutes incubation, 200 μL DNAse Stop Solution was added and samples were centrifuged for 30 seconds. Columns were washed with 600 μL RNA Wash Solution followed by 250 μL RNA Wash Solution, and then centrifuged again for 1 minute to dry. RNA was eluted using 100 μL of nuclease-free water. RNA quantification was performed using the Qubit RNA High Sensitivity Assay kit (Q32852).

### Quantitative Real-Time PCR (q-RT PCR)

To determine expression levels of *ompC/F*, *csgA/B* and *ramA,* we performed q-RT PCR using the Luna Universal One-Step RT-qPCR Kit from NEB (E3005), using the Applied Biosystems^ΤΜ^ 7500 Real-Time PCR system. The primers used for the q-RT PCR are listed in Table 2. Efficiency of the primers was calculated by generation of calibration curves for each primer pair on serially diluted DNA samples. The R^2^ of the calibration curves calibrated was ≥0.98 for all the primer pairs used in this study.

**Table 1.**
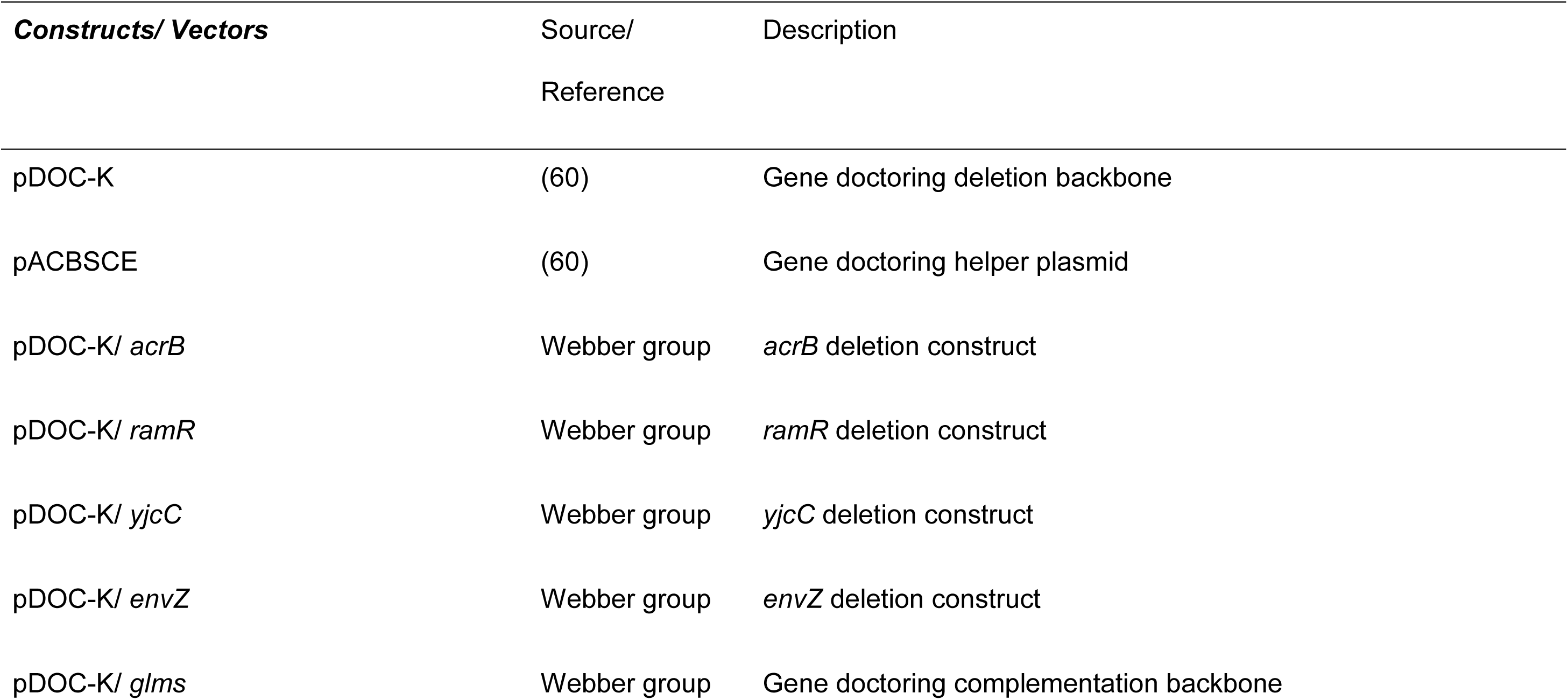
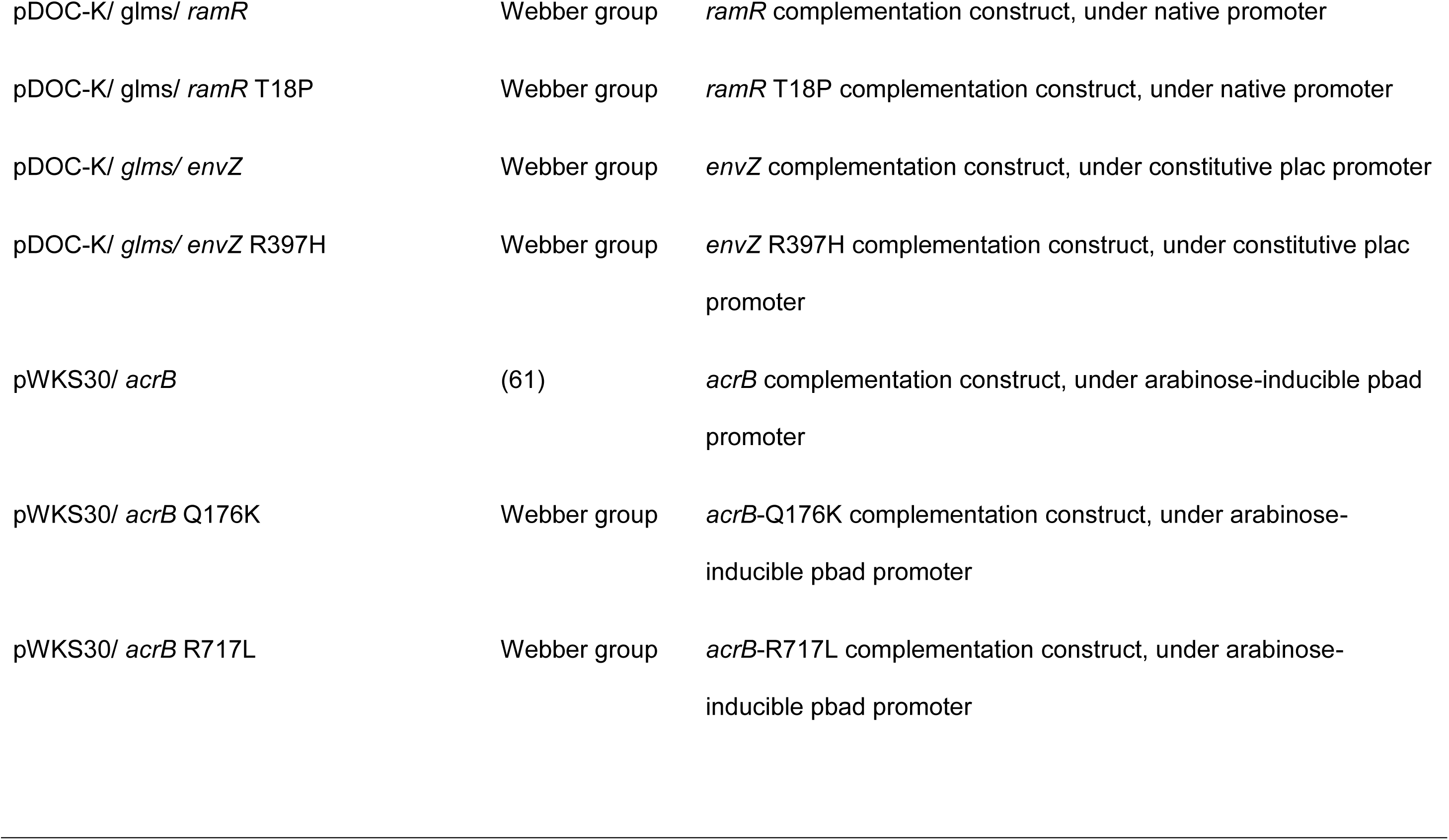
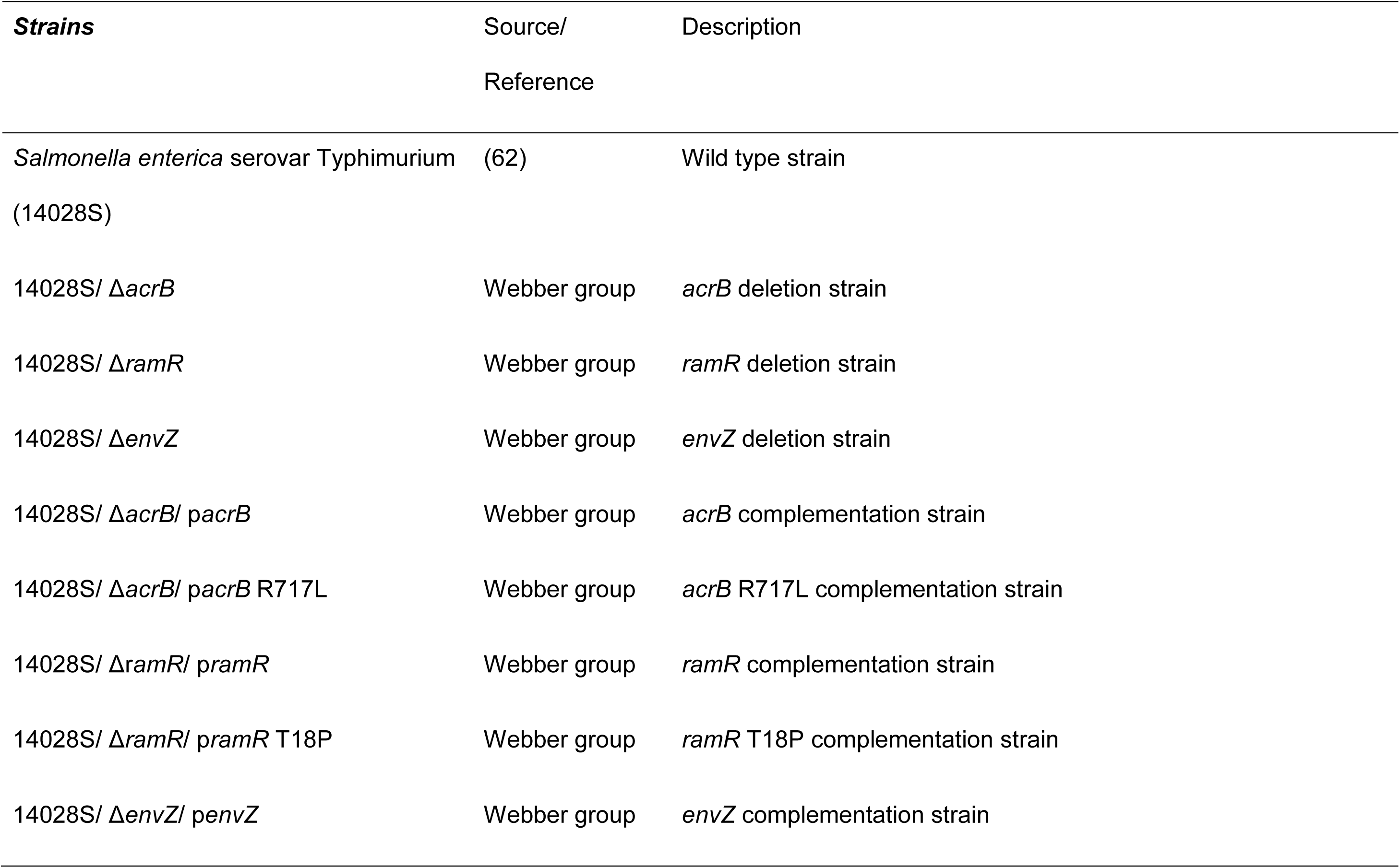
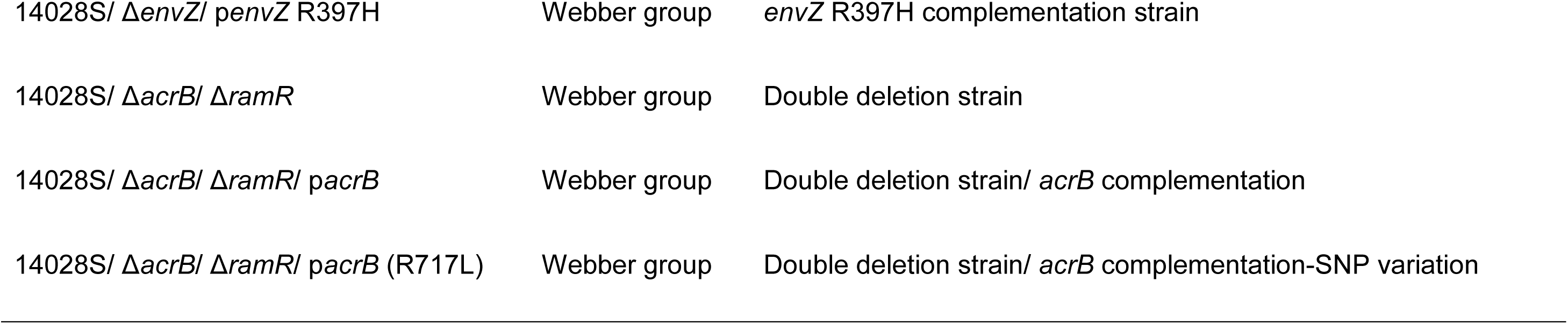
Bacterial strains and vectors.

**Table 2.**
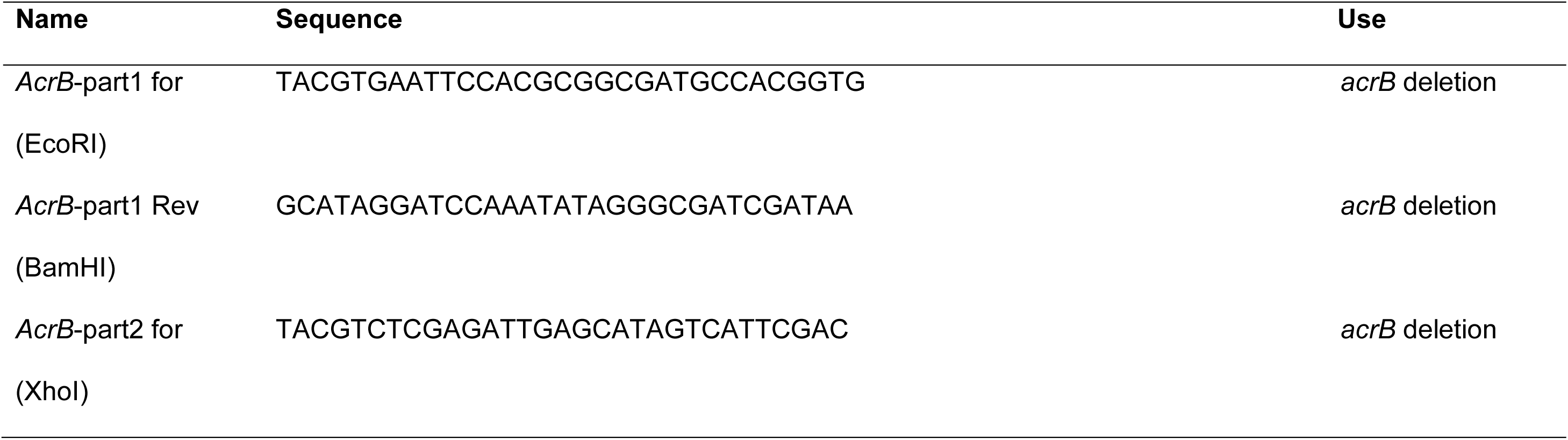
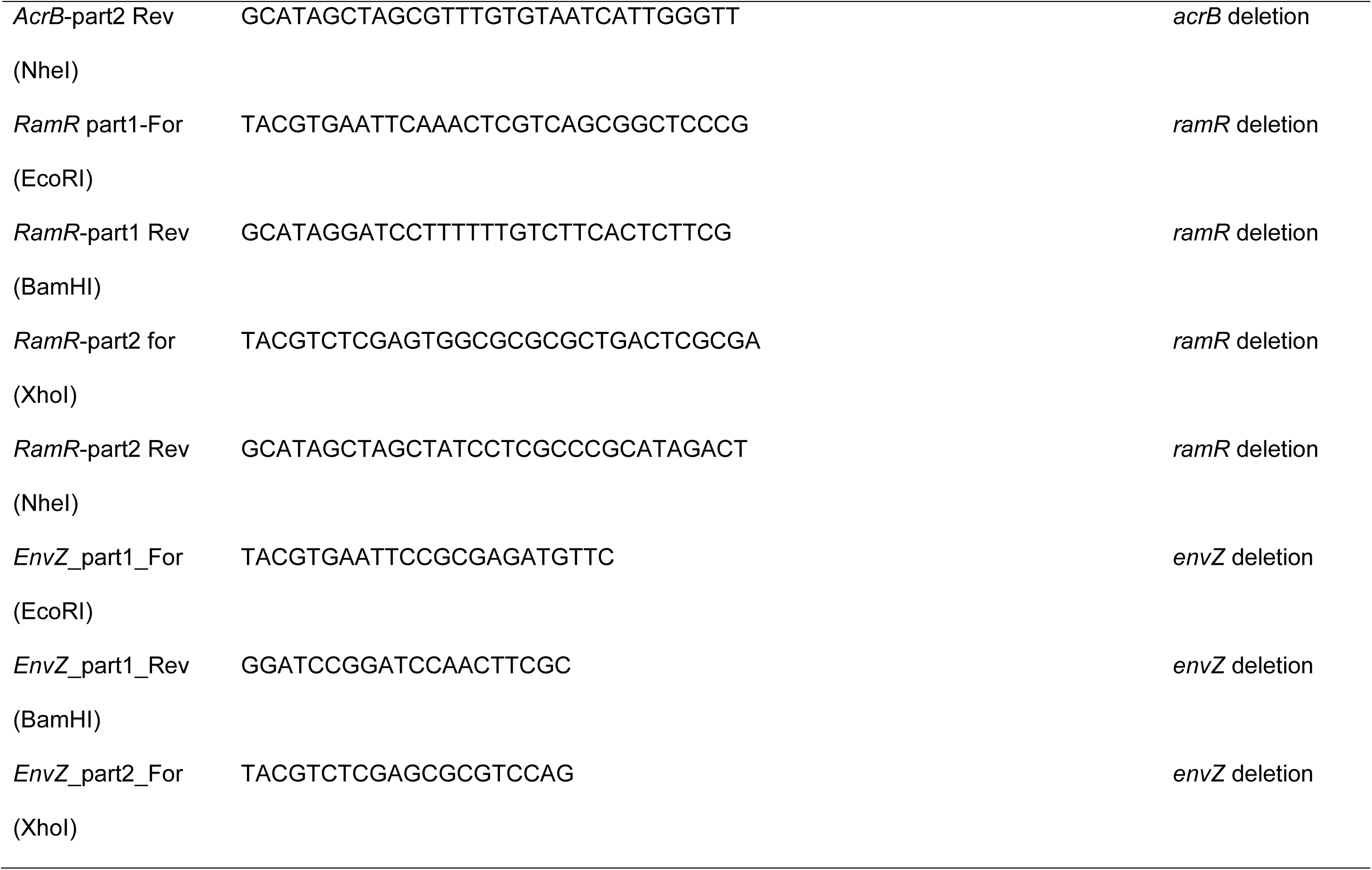
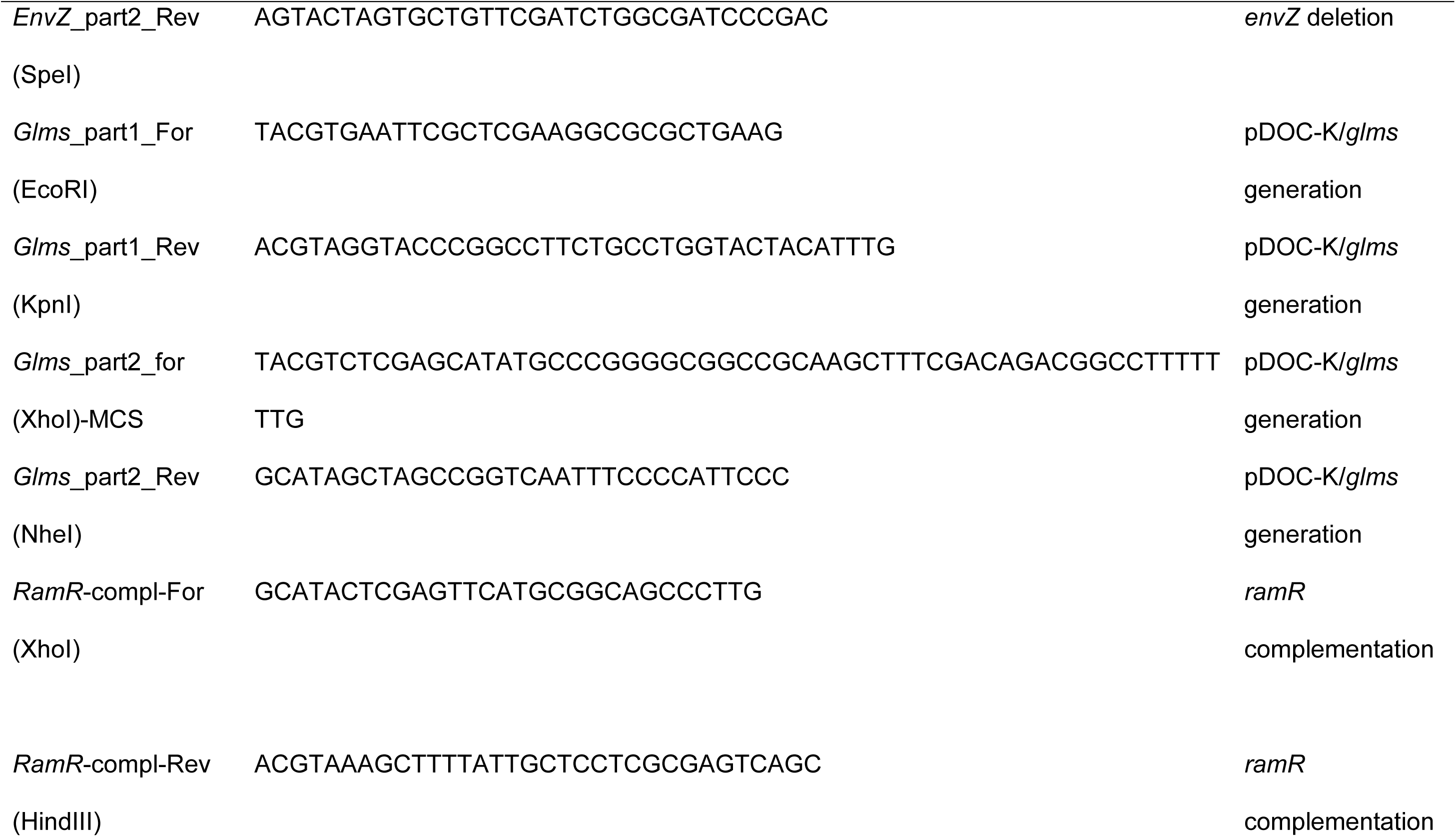
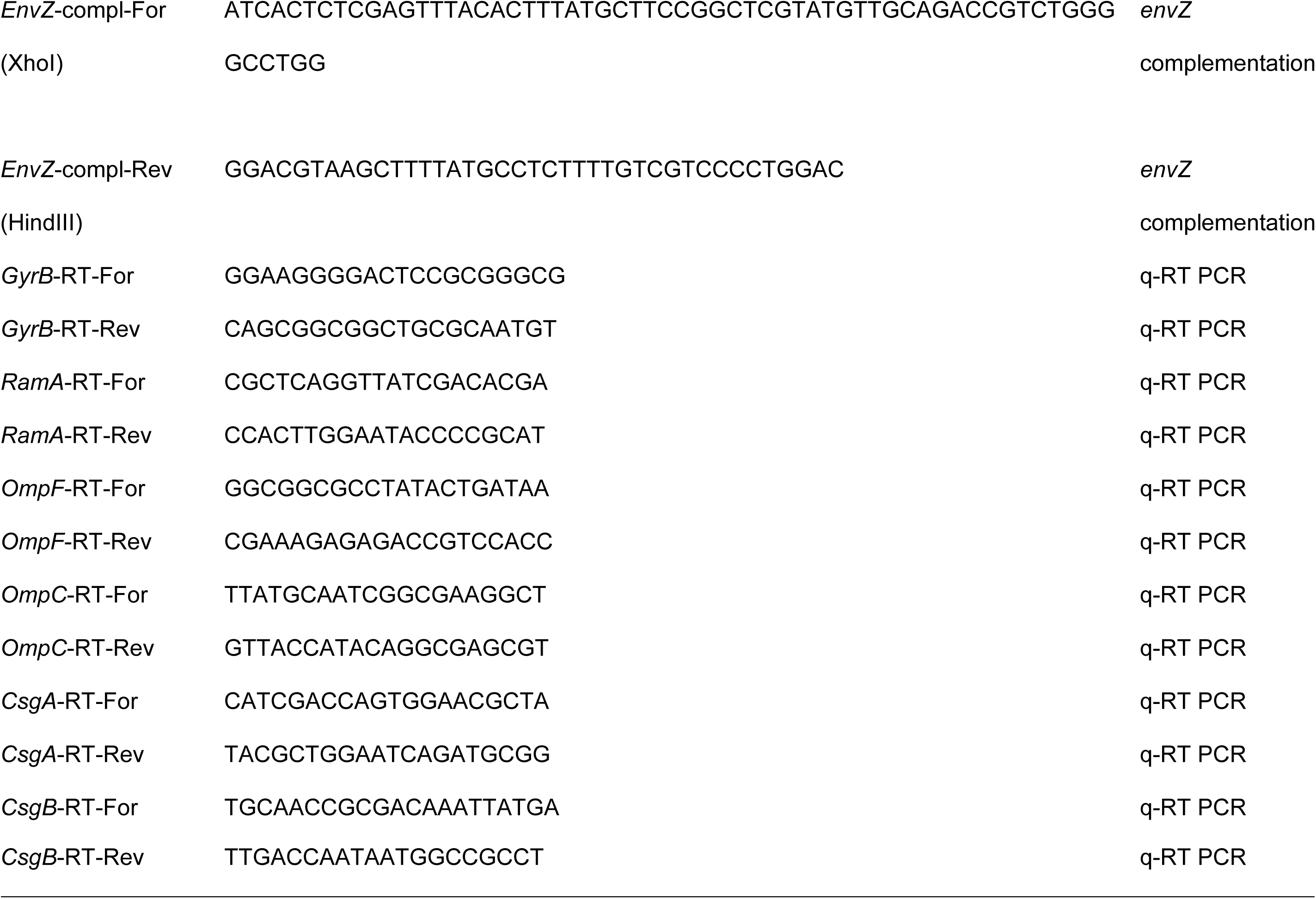
Primers used in this study.

RNA at a final amount of 50-100 ng was added to 10 μL final volume PCR reactions, mixed with 400 nM of each primer. The cycle parameters were as follows: 10 minutes at 55 °C (reverse transcription step), 1-minute denaturation at 95 °C and 40 cycles of 10 seconds at 95 °C and 1 minute at 60 °C.

For each sample, two technical replicates from two biological replicates each were included (four in total) per reaction. Controls with no reverse transcriptase were also included for each RNA sample to eliminate DNA contamination.

To calculate expression levels, expression fold change was calculated using *gyrB* expression as a reference. The relative expression was determined by calculating the logarithmic base 2 of the difference between *gyrB* gene expression and target gene expression per sample.

### Viability of cells within biofilms

To determine the viability of cells within a biofilm, two approaches were used. The first approach involved growing biofilms on glass beads for 72 hours. They were washed in PBS to remove planktonic cells and were then challenged with different concentrations of ciprofloxacin (0, 0.03, 0.3, 3 μg/mL) for 90 minutes. Beads were washed again in PBS to remove any antibiotic and transferred into 1 mL of PBS solution to an Eppendorf tube, where they were vigorously vortexed for 1 minute. The cells recovered in PBS were serial diluted and spotted onto LB plates for CFU counting the next day. For the second approach, we grew biofilms on glass slides for 72 hours. The slides were washed in PBS and were challenged with ciprofloxacin (3 μg/mL) for 90 minutes. They were washed in PBS and stained with a solution of 12 μM propidium iodide (PI) and 300 nM of SYTO 9 for 30 minutes. They were washed in PBS and soaked in 70% ethanol to kill the cells before they were transferred to a slide for microscopy. Fluorescence microscopy was performed in a Zeiss Axio Imager M2.

### Galleria Infection model

To test the pathogenicity of different mutants, we used the *Galleria mellonella* larvae infection model. Wax worms were obtained from livefoods.co.uk. Similarly-sized larvae with no signs of pupation or melanisation were chosen for injection. An initial experiment was performed to calculate the infectious dose of *S.* Typhimurium 14028S in *G. mellonella*, which determined that an inoculation with approximately 20,000 CFU resulted in death of approximately half of 10 larvae after 72 hours. Once this had been determined, overnight cultures of each strain were diluted in PBS to replicate this inoculum concentration and 10 μL of this were injected into the second hindmost left proleg of ten larvae. To check the concentration of each inoculum, 100 μL of each dilution were also plated onto LB agar and incubated overnight at 37°C. CFUs were counted the next day and the inoculum concentration was confirmed. Controls included in this experiment included larvae injected with PBS only and un-injected larvae. All larvae were incubated at 37 °C and were checked three times a day for 3 days to record the survival rate. The experiment was repeated on three independent occasions, with 10 larvae randomly allocated per strain in each experiment. Survival was calculated as the percentage of surviving larvae 48 hours after injection (Figure 6, e).

### Cellular permeability assays

To detect differences in cellular permeability to drugs between mutants, the resazurin accumulation assay was used. The strains of interest were grown to exponential phase, using a 1:100 inoculum from overnight cultures. The cells were washed and resuspended in PBS normalising for cell density and they were mixed with resazurin to a final volume of 100 μL (to give 10 μg/mL) in round-bottom microtiter plates. Fluorescence was measured in the Omega FLUOstar plate reader at excitation 544 nm and emission of 590 nm. Five replicates were included per strain and resazurin-only reactions were used as controls. The assay was repeated at least twice with reproducible results observed each time.

### Accelerated evolution experiments

To test the phenotypic stability of strains recovered from the initial evolution experiments, we performed an accelerated evolution experiment using six strains representing a spectrum of biofilm forming capacities and drug resistance phenotypes (WT, control-biofilm-L-S1, azi-biofilm-M-B-S2, azi-biofilm-M-B-S3, cef-biofilm-L-A-S1, cip-biofilm-M-B-S2, cip-biofilm-L-B-S3). The strains were resuscitated from storage by a 24-hour incubation at 37 °C in LB broth. After incubation, 50 µL of broth was added to 5 mL of LB broth (without salt) containing three sterile glass beads and incubated for 24 hours at 30 °C, until a biofilm was formed. Each bead was then washed in 1 mL PBS to remove planktonic and loosely adherent cells. Two beads were stored in deep-well plates containing 20 % glycerol for archiving and phenotyping. The third bead was transferred to another tube of LB broth (without salt) containing three sterile glass beads and passaged. This was repeated for ten passages, storing beads at each timepoint.

Upon completion of ten passages, populations were recovered from passage five, passage ten and the parental population for each mutant. From each population, single colonies were picked after streaking out each population on LB agar and incubating for 24 hours at 37 °C. Three colonies from each population were then subcultured in LB broth. A population and three isolates from the start, middle and end of the passage series were isolated and phenotyped for each mutant. Biofilm formation was evaluated using the Congo Red and Crystal Violet assays. The agar dilution methodology was used to assess the minimum inhibitory concentrations of antibiotics. The average of the fold MIC change per antibiotic for all strains was calculated and plotted against time. The average of biofilm formation, as determined by the crystal violet assay, was calculated for all the strains per timepoint.

### Statistical analysis

Biofilm forming ability was compared between strains or time points using linear mixed models, with a random intercept of lineage where more than one lineage was included for each strain or condition (Figures 1f and 6g).

Surviving cell counts were compared between strains using a linear mixed model with a Poisson response, with random intercept of replicate, fixed effects of exposure, the interaction between strain and exposure, and offset by the log of the average number of cells counted in the ‘unexposed’ condition for each strain. Modelled means in each exposure were then normalised by the average number of cells across all unexposed conditions for plotting, such that the values shown represent the estimated proportion of cells that would survive each exposure for each strain (Figure 6c). All error bars reflect estimates +- one standard error.

### Strains and genetic manipulations

*Escherichia coli* DH10b was used as a host for all cloning procedures. Transformations of *E. coli* were carried out by heat shock of chemically competent *E. coli* cells. Transformation of *Salmonella* was carried out by electroporation. *Salmonella* electrocompetent cells were prepared as follows: *Salmonella* cells were grown to early exponential phase (OD_600nm_ 0.2-0.3) in 50 mL of 2x YT, using a 1:100 inoculum from an overnight culture. The cells were centrifuged and washed once with filter sterilised ice-cold water. They were left to incubate on ice for 1 hour before they were pelleted at 3,000 g for 15 minutes. The cell pellet was resuspended in 1 mL of 10% filter-sterilised glycerol and 100 μL were used per transformation.

To create gene deletion mutants, we used the λ-red-based, gene doctoring technique previously described in (60). For each deletion, two homologous regions upstream and downstream of the genes of interest were amplified by PCR and were cloned in MCS1 and MCS2 of the pDOC-K vector. The homologous regions were 300-400 bp in length and were designed to include the first and last 10 codons of the gene to be deleted, to avoid any pleiotropic effects after deletion. For *acrB* and *ramR* deletions, the upstream homologous regions were cloned EcoRI/ BamHI in MCS1 and the downstream ones as XhoI/ NheI in MCS2 of pDOC-K. For the *envZ* deletion, the upstream homologous region was cloned EcoRI/ BamHI in MCS1 and the downstream one as XhoI/ SpeI in MCS2.

For complementation of mutated genes, chromosomal integrations were created to insert wild-type copies or mutated versions of genes of interest. This used a modification of the gene doctoring system described above. pDOC-K was modified to be used to deliver chromosomal gene integrations to the neutral intragenic region downstream of *glms*. This chromosomal insertion site was proven to be neutral through insertion of the reporter gene *lacZ*, where the mutant had no significant difference in biofilm formation, efflux activity or competitive fitness (data not shown). The integration vector (pDOC-K/ glms) was generated by cloning of the first homologous region in MCS1 of pDOC-K, using EcoRI/ KpnI and the second homologous region in MCS2 using XhoI/ NheI. The primer used for the amplification of the second homologous region was designed to introduce a novel MCS (XhoI, NdeI, SmaI, NotI, HindIII), upstream of the second homologous region which was then used for the gene complementation constructs. Wild-type *ramR* and ‘*ramR*-T18P’ alleles were cloned XhoI/ HindIII in pDOC-K/ glms under the control of the gene’s native promoter. Wild-type *envZ* and ‘*envZ-*SNP*’* alleles were cloned XhoI/ HindIII in pDOC-K/ glms under the control of a constitutive plac promoter. For *acrB* complementation, we used the pWKS30/ AcrB plasmid previously described in (61), expression of the gene is under the control of the pBAD system and induction was achieved with the use of 0.5% arabinose.

For induction of chromosomal integrations either for deletion or complementation of a gene, the strain to be modified was transformed with the pDOC-K vector variant and the pACBSCE helper plasmid carrying the λ-red genes. A single colony carrying both plasmids was grown in 500 μL of LB, at 37 °C for 4 hours. The cells were pelleted and washed three times in filter sterilized LB. They were then resuspended in 500 μL of 0.1x LB supplemented with 0.3% arabinose and incubated at 37 °C for 2-3 hours for induction. 100 μL were plated on LB plates supplemented with 25 μg/ mL kanamycin and 5% sucrose. The plates were incubated overnight at 37 °C. Single colonies were checked for chromosomal alterations using colony PCR with primers annealing outside of the region to be modified. The plasmids were removed by sub culturing the positive clones on kanamycin-supplemented plates and testing them for chloramphenicol and ampicillin sensitivity until the plasmids were completely removed.

For ‘double deletions’ and/or complementations, the kanamycin cassette, introduced by the first chromosomal modification, was removed using the FLP sites flanking the cassette. The strains were transformed by electroporation with the pCP20 vector, carrying the genes for flippase activity, and recovered on LB agar plates supplemented with 50 μg/mL ampicillin at 30 °C. The kanamycin cassette removal was confirmed by colony PCR and the positive clones were sub-cultured on LB agar at 37-42 °C. Removal of the plasmid was confirmed by testing the colonies’ sensitivity to ampicillin.

## Acknowledgements

We would like to thank David Baker and Gemma Kay for assistance with sequencing and Jacob Malone and Jessica Blair for helpful comments on the manuscript.

## Transparency declaration

The funders had no role in study design, data collection and analysis, decision to publish, or preparation of the manuscript

