## supplementary material for "Antibiotics select for novel pathways of resistance in biofilms"

**Supplementary Information - Docking**

For docking analysis, all target proteins and ligands were prepared in PDB format. Proteins and ligands PDB files were first converted to PDBqt format using Raccoon ^1^. AutoDock Vina ^2^ was used for docking simulation of ligands on protein structures (Table S1).

***Analysis of cefotaxime binding***

In the absence of experimental structures of cephalosporins bound to AcrB available, we selected chain B of 4DX5.pdb (minocycline coordinated in the distal binding pocket) as a template for the analysis of the effects of Q176K mutation ^3^. Chain B represents a drug-occupied conformation of the AcrB protomer. Comparing to previously published docking and MD data ^4^, 4DX5 was selected because of its higher resolution (1.9 Å versus 2.54 Å). For analysis of cefotaxime binding in the distal binding pocket, residues previously-characterised as involved in coordination of the ligand were selected (Q176, F178, I277, I278, F610, V612, F615, F617, F628) ^3,5,6^. These residues were defined as flexible during the docking, allowing for rotamer search. Docking scores are presented as free energy of binding (ΔG) (Table S1).

To validate our approach, we docked nitrocefin and cephalothin into chain B of 4DX5.pdb and in chain B of 2J8S.pdb. The latest has previously been used for the MD simulations of the above ligands ^4^. The top ligand binding poses are compared in Figure S1, a-b, demonstrating that the binding orientation is consistent between 4DX5.pdb and 2J8S.pdb, as well as with the previously published data ^4^.
Having shown the validity of the approach, we then performed further simulations to analyse the binding of cefotaxime into two separate templates based on the experimental 4DX5.pdb chain B structure, representing the wild type (Q176) and mutant binding pocket (Q176K) respectively (Figure 4, e-f main text and Figure S1, c). Importantly the Q176 residue was found to coordinate cefotaxime in the WT pose, while the Q176K substitution resulted in a notably different orientation and less favourable energy of binding (Table S1).

Furthermore, the orientation of the top pose of cefotaxime differed notably from the orientations of both cephalothin and nitrocefin in our simulations, in both WT and mutant proteins. The molecule appears to be flipped 180 degrees along its long axis, and, while Q176 is involved in the binding of nitrocefin, the coordination of cefotaxime is not comparable (Figure S1, d). This can be explained by the significantly different physicochemical properties of the compound.

***Analysis of Azithromycin binding***

Previous crystallographic evidence ^7^, has shown high molecular weight drugs (HMMD) bind in a partially-overlapping multisite binding pocket (aka proximal binding pocket) of the "binding" or "access" protomer of AcrB. In the case of the macrolides, two distinct sub-pockets have been designated as site A and B, corresponding to the sites occupied by rifampicin and erythromycin in the structures 3AOB, chain C and 3AOC, chain C respectively. While azithromycin is closely related to erythromycin, we tested binding to both sites, as site A forms a logical access path to site B and may be at least partially responsible for the vetting of the ligand.

***Analysis of azithromycin binding to macrolide site B***

For the analysis of the effect of R717L on macrolide site B of the proximal pocket of AcrB, chain C from the AcrB-erythromycin co-crystal structure (3AOC.pdb) was used. For analysis of erythromycin and azithromycin binding in the proximal binding pocket, residues previously-characterised as involved in coordination of the ligand were selected (S79, Q81, S134, F136, K292, M575, E673, D681, F617, R717, F666, E826) ^5,7^. These residues were defined as flexible during the docking, allowing for rotamer search. The simulation was run with both WT R717 and the R717L mutant. Docking scores are presented as free energy of binding (ΔG) (Table S1).

Using a similar approach to the cefotaxime binding study described above, to validate the model, erythromycin and azithromycin were docked to macrolide site B of the proximal binding pocket and compared with the experimental erythromycin-containing structure 3AOC, chain C ^7^ (Figure S2, a-b).

***Analysis of azithromycin binding to macrolide site A***

For the analysis of the macrolide binding to the macrolide site A of the proximal pocket, a template was designed based on chain C of 3AOB.pdb, corresponding to the AcrB–rifampicin complex. Residues previously-characterised as involved in coordination of the ligand were selected as flexible (S134, F136, Q577, M575, M573, N623, T624, M662, L674, F617, R717, N719, F664, F666, R815, L828) ^7^. Azithromycin was docked to templates containing both R717 WT and R717L substitution (main text Fig. 5 G-H); Rifampicin and erythromycin were docked as controls and the resulting poses were comparable to the experimental rifampicin-containing structure (3AOB, chain C) (Figure S2, c-d).

**Figures**

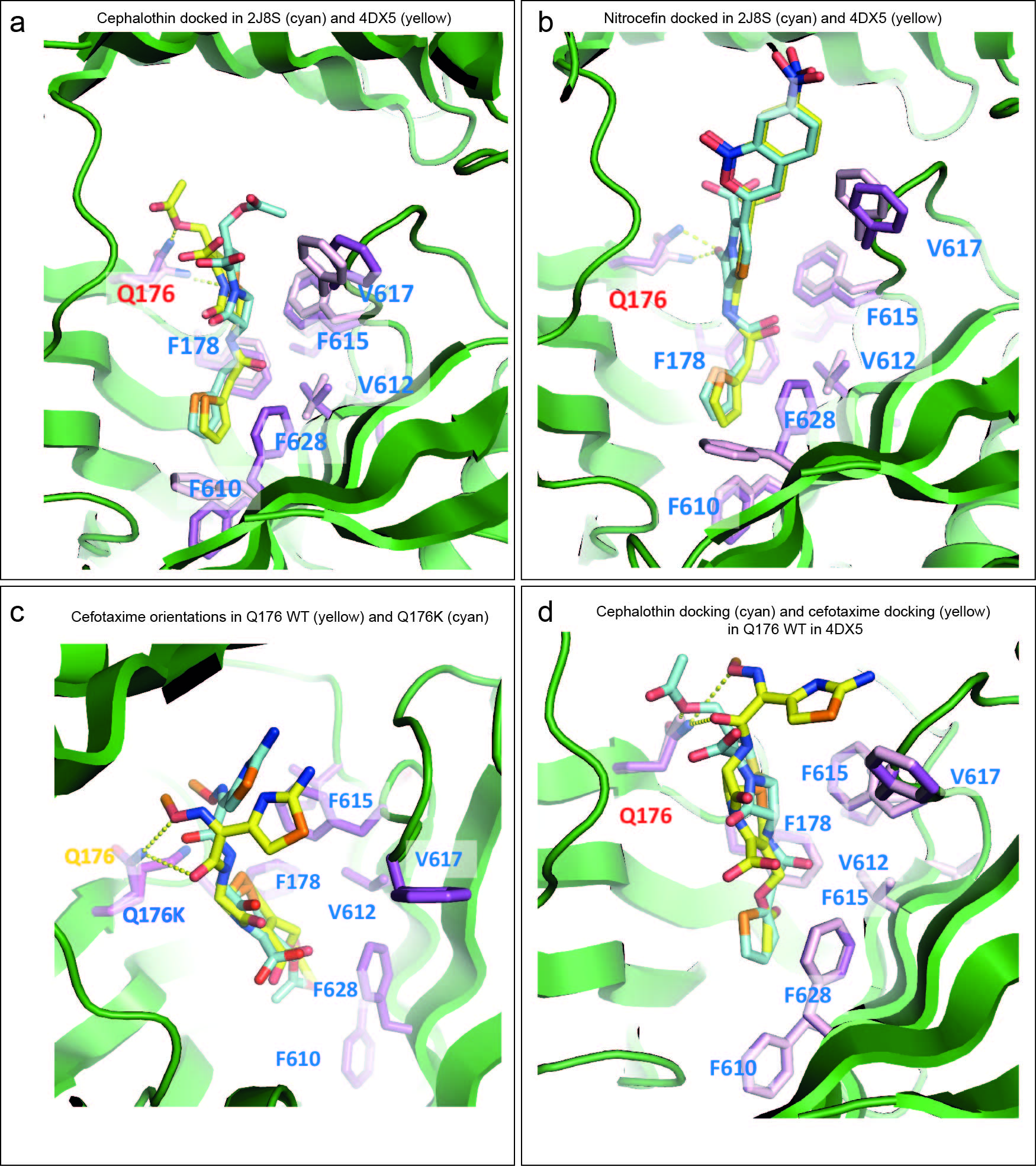
Figure S1

***Figure S1. Cefotaxime docking validation. a,*** *Comparison between the docked poses of cephalothin in 2J8S (cyan) and 4DX5 (yellow) reveals consistent orientation of the ligand.* ***b,*** *Comparison between the docked poses of nitrocefin in 2J8S (cyan) and 4DX5 (yellow) reveals consistent orientation of the ligand.* ***c,*** *Comparison of the orientations of the top cefotaxime poses for Q176 WT (yellow) and Q176K (cyan). The position of the mutated side chain differs significantly between the two pockets.* ***d,*** *Comparison of the orientations of the top cefotaxime pose (yellow) with the top cephalothin pose (cyan) docked to the WT protein. The long axis of cefotaxime is flipped by 180^o^, in comparison with the cephalothin and nitrocefin structures (a and b), due to different charge distribution.*

Figure S2

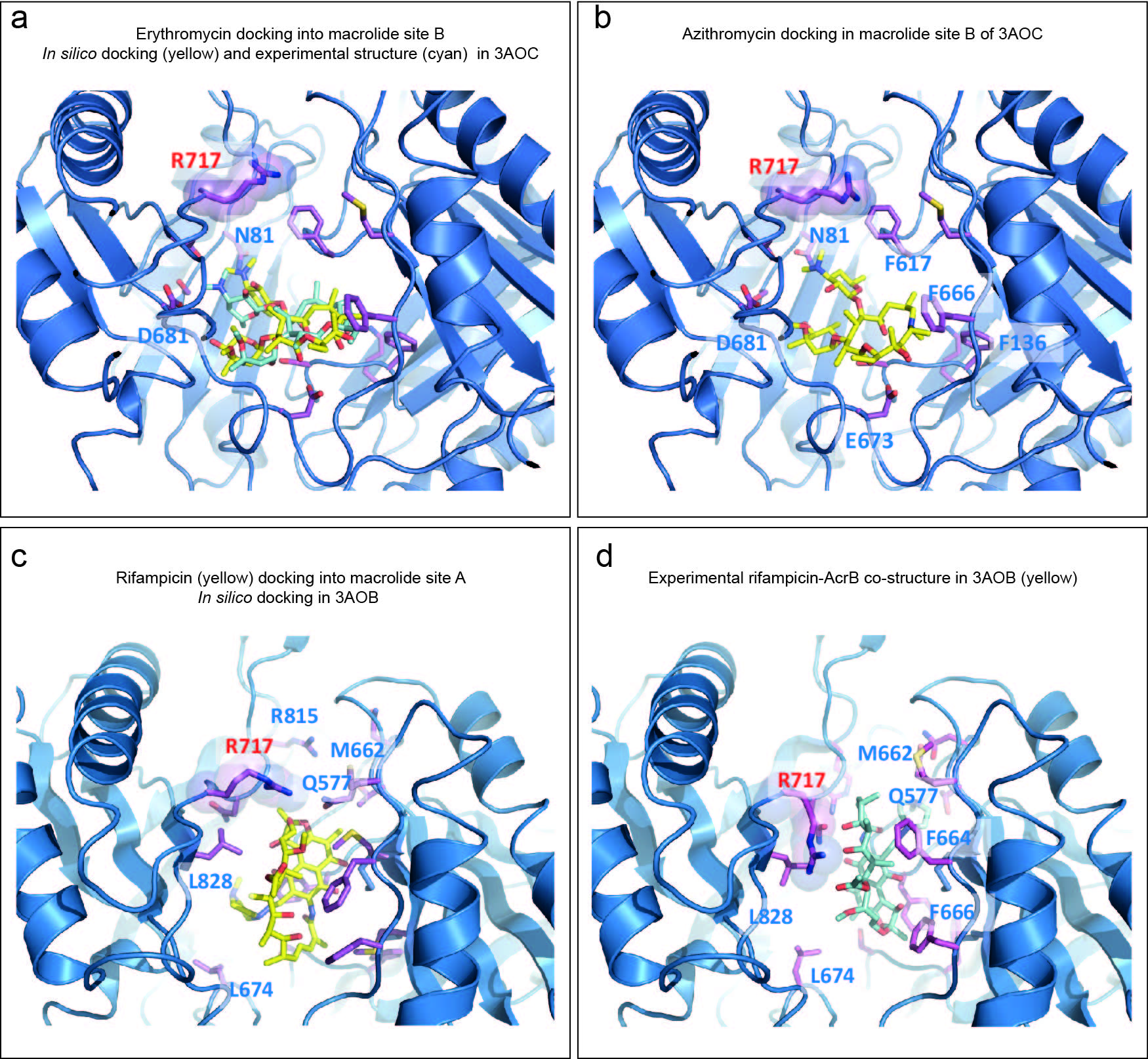

***Figure S2. Azithromycin docking validation****.* ***a,*** *Erythromycin in silico docking to macrolide site B (yellow) and comparison with the experimental crystal structure 3AOC (cyan).* ***b,*** *Azithromycin docking into macrolide site B reveals that the position is highly consistent with that of erythromycin and presents a conservative interaction. However, R717 remains too far for coordination to be possible.* ***c,*** *Best energy pose for in silico docked rifampicin into 3AOB binding pocket (flexi-dock) with flexible rotamers in sticks. R717 makes clear contact with the ligand.* ***d,*** *Experimental co-crystal structure of AcrB with rifampicin (3AOB).*

Table S1. Docking energies

| Affinity of binding (Kcal/mol) | | |
| --- | --- | --- |
| **Distal binding pocket**  **Docking box (Å): 24x28x22** | | |
| **Protein (AcrB)** | **WT (Q176)** | **WT (Q176)** |
| **Ligands/ Structures (pdb)** | **4DX5** | **2J8S** |
| Cephalothin | -9.5 | -9.1 |
| Nitrocefin | -11.1 | -10.8 |
| **Protein (AcrB)** | **WT (Q176)** | **Q176K** |
| **Ligands/ Structures** | 4DX5 | 4DX5 |
| Cefotaxime | -8.4 | **-8.0** |
| **Proximal binding pocket**  **Docking box (Å): 26x26x22** | | |
| **Protein (AcrB)** | **WT (R717)** | |
| **Ligands/ Structures (pdb)** | **3AOC** | |
| Azithromycin | -12 | |
| Erythromycin | -12.7 | |
| **Docking Box (Å): 32 x 30 x 30** | | |
| **Protein (AcrB)** | **WT (R717)** | **R717L** |
| **Ligands/ Structures (pdb)** | **3AOB** | **3AOB** |
| Azithromycin | -11.2 | **-10.3** |
| Erythromycin | -11.0 | **-10.4** |
| Rifampicin | -11.9 | -11.6 |
